# Distinct Roles of Dopamine D1 and D2 Receptor-expressing Neurons in the Nucleus Accumbens for a Strategy Dependent Decision Making

**DOI:** 10.1101/2021.08.05.455353

**Authors:** Tadaaki Nishioka, Tom Macpherson, Kosuke Hamaguchi, Takatoshi Hikida

**Author notes:** Correspondence (T.N.), (T.H.).

## Abstract

To optimize decision making, animals need to execute not only a strategy to choose a good option but sometimes also one to avoid a bad option. A psychological study indicates that positive and negative information is processed in a different manner in the brain. The nucleus accumbens (NAc) contains two different types of neurons, dopamine D1 and D2 receptor-expressing neurons which are implicated in reward-based decision making and aversive learning. However, little is known about the neural mechanisms by which D1 or D2 receptor-expressing neurons in the NAc contribute to the execution of the strategy to choose a good option or one to avoid a bad option under decision making. Here, we have developed two novel visual discrimination tasks for mice to assess the strategy to choose a good option and one to avoid a bad option. By chemogenetically suppressing the subpopulation of the NAc neurons, we have shown that dopamine D2 receptor-expressing neurons in the NAc selectively contribute to the strategy to avoid a bad option under reward-based decision making. Furthermore, our optogenetic and calcium imaging experiments indicate that dopamine D2 receptor-expressing neurons are activated by error choices and the activation following an error plays an important role in optimizing the strategy in the next trial. Our findings suggest that the activation of D2 receptor-expressing neurons by error choices through learning enables animals to execute the appropriate strategy.

## INTRODUCTION

Even seemingly similar decisions could be mediated by different cognitive processes. For example, selecting a food for dinner could be mediated by either picking up one’s favorite food or avoiding the one with a bad experience. Although selecting a better choice and avoiding a bad choice could lead to a similar choice behavior, a psychological study indicates that their underlying processes in the brain could be different (Tversky and Kahneman, 1981). A number of studies have proposed the basal ganglia circuitry is implicated in decision making (Cohen et al., 2012; Frank et al., 2004; Hamid et al., 2015; Hikida et al., 2010; Iino et al., 2020; Maia and Frank, 2011; Samejima et al., 2005; Schultz et al., 1997; Stephenson-Jones et al., 2016; Tai et al., 2012; Yagishita et al., 2014; Zalocusky et al., 2016). Dopamine has been known as an important modulator of basal ganglia functions (Ferenczi et al., 2016; Gerfen and Surmeier, 2011; Parker et al., 2018; Shen et al., 2008; Surmeier et al., 2009). The nucleus accumbens (NAc), one of the major projection targets of dopamine neurons, is known to play an important role in reward-based decision making and aversive learning (Berridge and Kringelbach, 2008; Carlezon and Thomas, 2009; Hikida et al., 2010; Iino et al., 2020; Lee, 2013; Yawata et al., 2012; Zalocusky et al., 2016). The medium spiny projection neurons (MSN) in the NAc are divided into two major subpopulations, one that expresses the dopamine D1 receptor (D1-MSN) and the other that expresses the dopamine D2 receptor (D2-MSN). Because D1 and D2 receptors have antagonistic response to dopamine, D1- and D2-MSN were thought to have different contribution to the choice behavior. Previous studies showed that D1-MSN and D2-MSN in the NAc play distinct roles in a context dependent manner (Aragona et al., 2006; Hikida et al., 2010; Lobo et al., 2010; Macpherson and Hikida, 2018; Yawata et al., 2012), however, it is not well understood how and whether different strategies, such as selecting a better choice or avoiding a bad choice, are differentially represented in these two distinct classes of neurons in NAc.

In this study, we have developed a pair of visual discrimination learning tasks to assess the strategy dependent decision making in mice. While in the visual discrimination-based cue-guided attendance learning (VD-Attend), mice need to acquire the strategy to choose a good option. In contrast, in the visual discrimination-based cue-guided avoidance learning (VD-Avoid), mice need to acquire the strategy to avoid a bad option. To establish the contribution of these two subpopulations in these different tasks, we chemogenetically manipulated the neuronal activities of D1-MSN or D2-MSN in the NAc while mice performing the VD-Attend or VD-Avoid tasks. Moreover, we performed optogenetic manipulation to test whether the NAc contributes to the execution of the strategy in a timing-specific manner. We further performed in vivo calcium imaging with a miniature microscope to investigate the underlying neural representations of D1-MSN and D2-MSN. Our results show that the majority of D1-MSN in the NAc is recruited by correct choice and the activity of D1-MSN is important for the motivational control and keeping the same strategy. By contrast, D2-MSN in the NAc plays a selective role in executing the strategy to avoid a bad option in VD-Avoid task. Furthermore, we found that the vast majority of D2-MSN are activated by error choices and the activation of D2-MSN after error is necessary for optimizing the strategy in next trial. Our findings suggest that the activation of D2-MSN by error choices through learning enables animals to execute the appropriate strategy.

## RESULTS

### Mice Can Acquire Both the Strategy to Choose a Good Option and One to Avoid a Bad Option

To study the neural mechanisms underlying the strategy to choose a good option and one to avoid a bad option, we designed two novel visual discrimination learning tasks, VD-Attend and VD-Avoid task on a touchscreen-based operant chamber (**Figure 1A**). In these tasks, two visual cues were presented on a touchscreen after trial initiation (**Figure 1B**). If mice responded to a correct cue, the reward was delivered in the magazine. If mice responded to an incorrect cue, they were punished with a 5-sec time-out. In the VD-Attend task, while the visual cue A (e.g. marble) was presented as a correct cue in every trial, a random image was presented as an incorrect cue. Therefore, mice were required to execute a strategy to choose cue A in order to obtain a reward (**Figure 1A**). By contrast, in the VD-Avoid task, the visual cue B (e.g. fan) was presented as an incorrect cue and a random image was presented as a correct cue. Therefore, mice cannot directly learn the correct cue, but instead, mice could obtain a reward by avoiding the cue B (**Figure 1A**). Next, we trained C57BL/6J mice to the VD-Attend or VD-Avoid task and confirmed that mice can perform both tasks well within a few weeks of training (**Figure 1C**). The number of sessions and total errors to criterion in the VD-Avoid task were larger than that in the VD-Attend task (**Figure 1D** and **1E**), suggesting that acquisition of the strategy to avoid a bad option is more difficult than that to choose a good option. However, the performances measured by the percentage of correct choice were comparable in the VD-Attend and VD-Avoid tasks across sessions (**Figure 1F and 1G**). Correct latency and reward latency were also comparable in the VD-Attend and VD-Avoid (**Figure 1H and 1I**). A number of studies have reported the phenomenon of post-error slowing in which the subjects tend to slow down the response after an error trial (Houtman et al., 2012; Notebaert et al., 2009; Rabbitt and Rodgers, 1977; Ruitenberg et al., 2014). We also tested whether the recent history on the previous trial affected the response latency on the current trial. We found that mice tended to respond slowly on the next trial after making an error response in the VD-Avoid task (**Figure S1**), suggesting that response latencies were modulated by recent history. Together, these data show that mice can acquire both strategies of attending and avoiding specific visual cues in the touchscreen-based operant tasks, which provide a framework for studying the neural mechanisms underlying two different forms of strategies.

**Figure 1.**
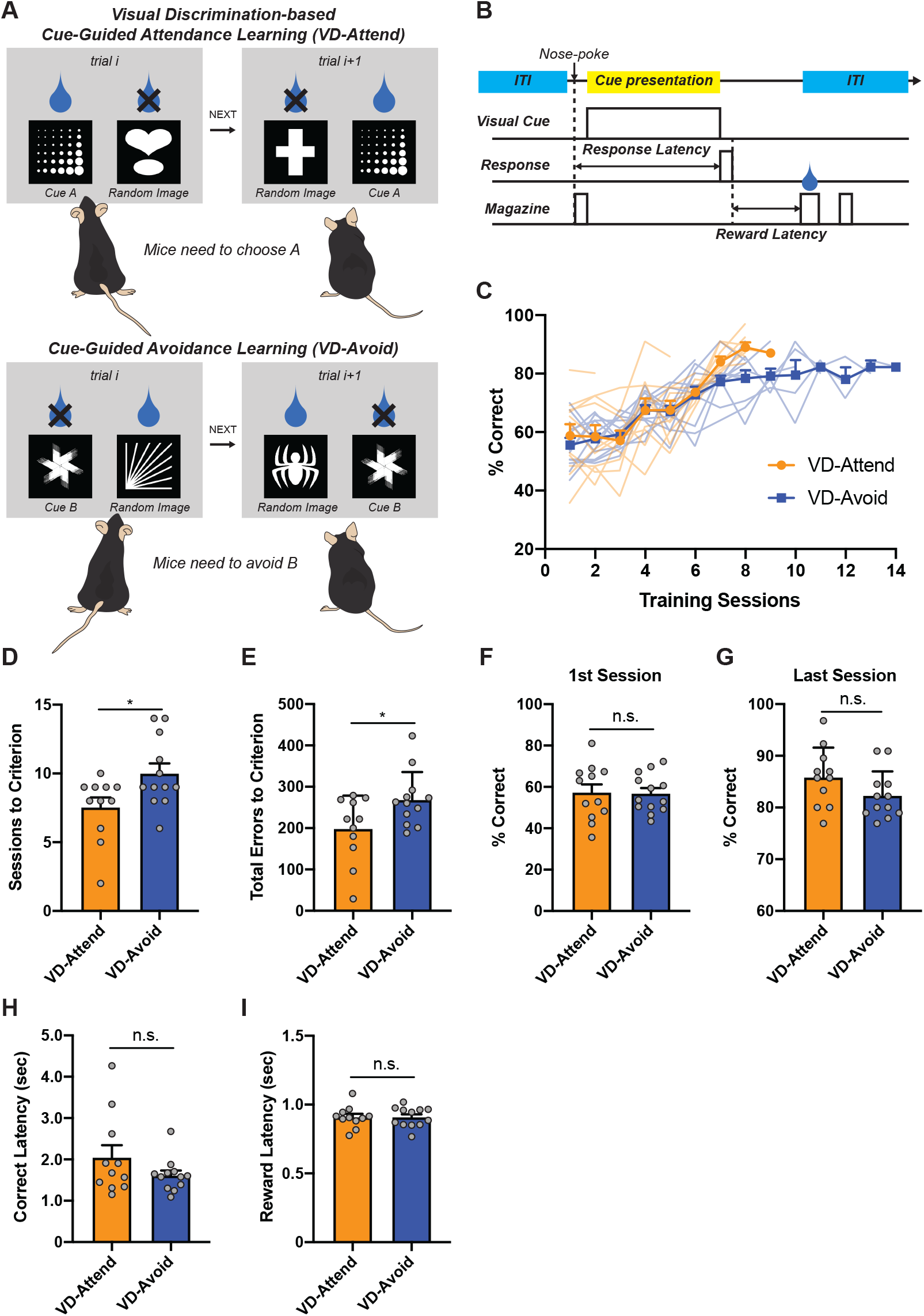
Experimental Design and Behavioral Performances. (A) Experimental design. (B) Timeline of the task events and the definition of the behavioral parameters. (C) The percentage of correct in each session (VD-Attend, n = 11; VD-Avoid, n = 12). (D) The number of sessions to criterion of the VD-Avoid was higher than that of the VD-Attend task (unpaired t-test, t_21_ = 2.413, *p = 0.0251). (E) Total errors to criterion of the VD-Avoid were higher than that of the VD-Attend task (unpaired t-test, t_21_ = 2.288, *p = 0.0326). (F and G) The behavioral performances were similar between the VD-Attend and VD-Avoid task in the 1^st^ session (F, unpaired t-test, t_21_ = 0.7053, p = 0.4884) and the last session (G, unpaired t-test, t_21_ = 1.613, p = 0.1217). (H and I) Correct latencies were similar between the VD-Attend and VD-Avoid task in the last session (H, unpaired t-test, t_21_ = 1.420, p = 0.1704). Reward latencies were similar between the VD-Attend and VD-Avoid task in the last session (I, unpaired t-test, t_21_ = 0.007948, p = 0.9937).

### Chemogenetic Suppression of D2-MSN in the NAc Selectively Impair the Ability to Execute the Strategy to Avoid a Bad Option

Given that dopaminergic medication impaired the ability to avoid a bad option option (Frank et al., 2004, 2007), and the NAc, one of the major targets of the dopamine neurons, plays an important role in decision making (Berridge and Kringelbach, 2008; Carlezon and Thomas, 2009; Hikida et al., 2010; Iino et al., 2020; Lee, 2013; Yawata et al., 2012; Zalocusky et al., 2016), we first investigated whether D1-MSN and D2-MSN in the NAc contribute to the performance in VD-Attend and VD-Avoid tasks. We bilaterally injected an adeno-associated virus (AAV) carrying inhibitory designer receptor exclusively activated by a designer drug (iDREADD, hM4Di) fused to mCherry (Armbruster et al., 2007) in a Cre dependent manner into the NAc of D1-Cre or D2-Cre mice (**Figure 2A**). The hM4Di can robustly suppress the neuronal activities via cell signaling when an artificial ligand clozapine-N-oxide (CNO) is administered (Wulff and Arenkiel, 2012). To test whether D1-MSN or D2-MSN in the NAc is involved in executing the strategy, we trained mice expressing hM4Di-mCherry in D1-MSN or D2-MSN of the NAc to the VD-Attend or VD-Avoid task (**Figure 2A**). After mice were well-trained, we intraperitoneally administered CNO at the session 4 and 6 in probe test (**Figure 2A**). In this study, we mainly targeted the core region of the NAc because recent studies indicate that the dopamine neurons are heterogeneous and the dopamine neurons projecting to the core but not shell region encode the reward prediction error information (Iino et al., 2020; Jong et al., 2019; Parker et al., 2016; Roitman et al., 2008). We histologically confirmed that hM4Di-mCherry expression in the injection site were mainly restricted to the NAc (**Figure 2B**). Moreover, in D1-Cre mice, hM4Di-mCherry-positive axon terminals were observed in the ventral pallidum (VP) and substantia nigra pars reticulata (SNr), while in D2-Cre mice, hM4Di-mCherry-positive axon terminals were observed only in the VP but not SNr (**Figure S2**), which are consistent with the canonical targets of D1-MSN or D2-MSN in the NAc (Lu et al., 1997; Zhou et al., 2003). These data demonstrates that the viral transduction allowed us to selectively express hM4Di-mCherry in D1-MSN or D2-MSN in the NAc. A previous study also shows that CNO can functionally suppress the neuronal activity of the NAc (Luo et al., 2018). Next, we examined how chemogenetic suppression of D1-MSN or D2-MSN in the NAc via CNO influenced the performance of the VD-Attend or VD-Avoid task across sessions (**Figure 2C**). Chemogenetic suppression of D1-MSN in the NAc reversibly decreased the performance of both the VD-Attend and VD-Avoid task (**Figure 2D**). Interestingly, chemogenetic suppression of D2-MSN in the NAc selectively decreased the performance of the VD-Avoid but not VD-Attend task (**Figure 2D**), suggesting that the neuronal activities of D2-MSN in the NAc is necessary for avoiding a bad option but not choosing a good option under decision. The extent of decreased performance in D1-Cre mice was comparable to that in D2-Cre mice in the VD-Avoid task (**Figure 2E**). Moreover, we confirmed that CNO itself did not affect the performance of the VD-Attend or VD-Avoid task though CNO treatment slightly improved the performance of the VD-Avoid task in control group (**Figure 2D**). Considering that the NAc plays an important role in motivational control (Aberman et al., 1998; Berridge, 2007; Delgado, 2007; Gallo et al., 2018; Roitman et al., 2005; Salamone et al., 2007; Schultz et al., 1992) and that a decrease in the performance in the reward-based decision making could be associated with motivation deficits, we analyzed the effect of chemogenetic manipulation on motivation. We found that chemogenetic suppression of D1-MSN in the NAc decreased the number of earned rewards and increased reward latency, both of which indicated decreased motivation (**Figure 2F and 2G**). On the other hand, chemogenetic suppression of D2-MSN in the NAc did not affect the motivation indices (**Figure 2F and 2G**), suggesting that a decrease in performance of the VD-Avoid task are attributed to the impaired ability to avoid a bad option. Furthermore, we analyzed whether the outcome of the previous trial affected the performance of the current trial. The chemogenetic suppression of D1-MSN in the NAc resulted in a decrease of performance following a correct trial (**Figure S3A and S3B**), suggesting that D1-MSN in the NAc plays an important role in consecutively making a correct choice. On the other hand, we could not detect the effect of the recent history on the behavioral performance in D2-Cre or control group (**Figure S3C-S3F**). Collectively, our data indicate that D1-MSN and D2-MSN in the NAc contribute to the choice behavior differently; D1-MSN contributes to both the VD-Attend and VD-Avoid tasks, and D2-MSN selectively involved in the VD-avoid task.

**Figure 2.**
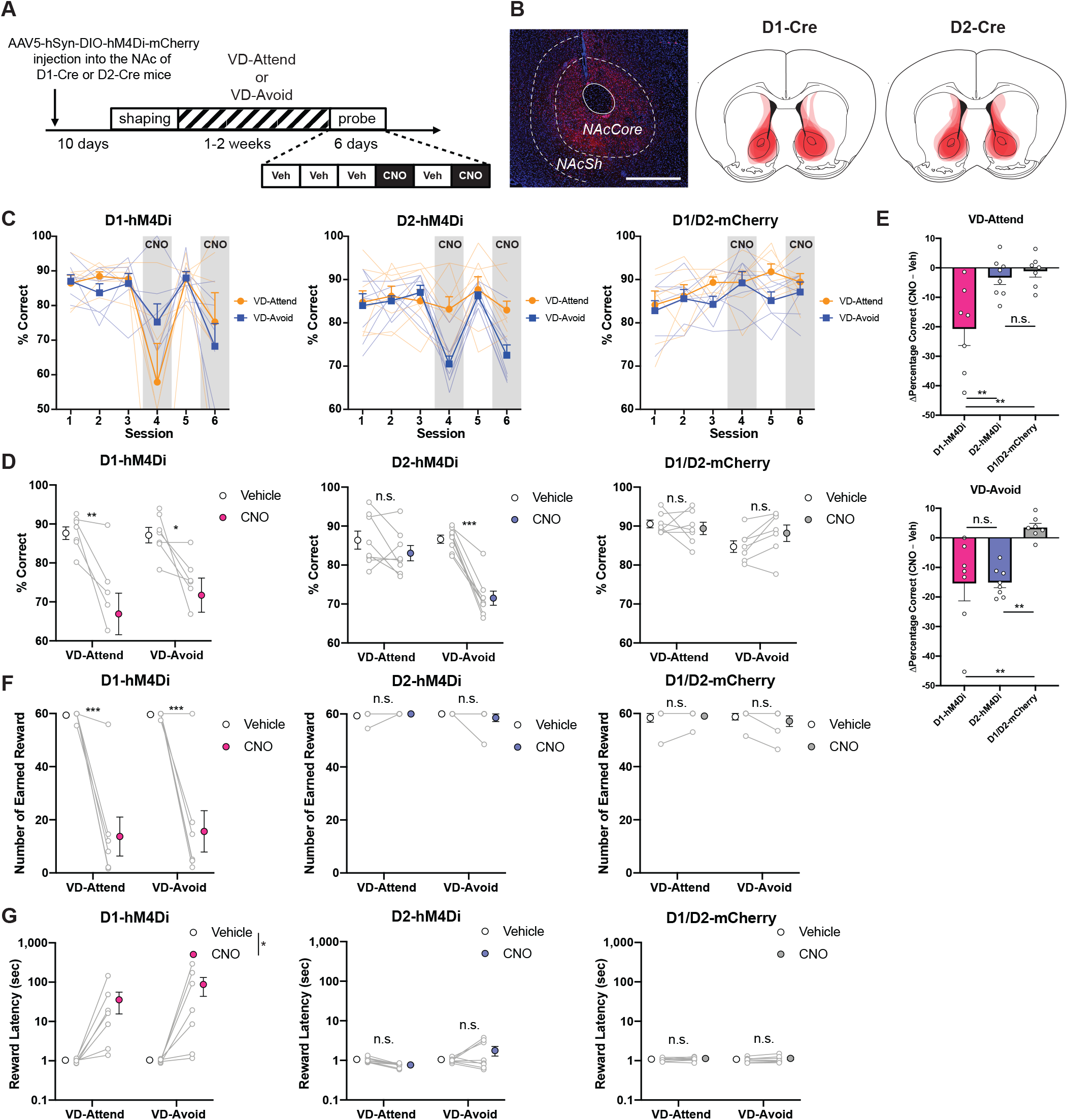
DREADD Suppression of D2-MSN in the NAc Selectively Decreases the Performance of the VD-Avoid Task. (A) Experimental timeline. (B) Representative injection site of AAV5-hSyn-DIO-hM4Di-mCherry (Left). Scale bar, 500 µm. NAcCore, nucleus accumbens core; NAcSh, nucleus accumbens shell. Drawings of superimposed AAV injection sites in the NAc (Right). (C) The effects of DREADD suppression on the behavioral performance of each task in D1-Cre mice (Left, n = 7), D2-Cre mice (Middle, n = 8). The effects of CNO treatment on the behavioral performance of each task in D1/D2-mCherry mice (Right, n = 7). (D) DREADD suppression of D1-MSN in the NAc decreased the behavioral performance in both the VD-Attend and VD-Avoid task (Left, Two-way RM-ANOVA with Sidak correction, Treatment effects, F_1,12_ = 19.83, ***p = 0.0008; VD-Attend, **p = 0.0071; VD-Avoid, *p = 0.0394). DREADD suppression of D2-MSN in the NAc selectively decreased the behavioral performance in the VD-Avoid but not VD-Attend task (Middle, Two-way RM-ANOVA with Sidak correction, Treatment effects, F_1,14_ = 40.88, ***p < 0.0001; VD-Attend, p = 0.2334; VD-Avoid, ***p < 0.0001). CNO treatment in D1/D2-mCherry did not affect the behavioral performance in both the VD-Attend and VD-Avoid task (Right, Two-way RM-ANOVA with Sidak correction, Treatment effects, F_1,12_ = 0.8707, p = 0.3692; VD-Attend, p = 0.7590; VD-Avoid, p = 0.1326). (E) The effects of DREADD suppression on the behavioral performance of the VD-Attend task across all groups (Top, One-way ANOVA with Sidak correction, Group effects, F_2,19_ = 8.688, **p = 0.0021; D1-hM4Di vs D2-hM4Di, **p = 0.0078; D1-hM4Di vs D1/D2-mCherry, **p = 0.0039; D2-hM4Di vs D1/D2-mCherry, p = 0.9643). The effects of DREADD suppression on the behavioral performance of the VD-Avoid task across all groups (Bottom, One-way ANOVA with Sidak correction, Group effects, F_2,19_ = 9.396, **p = 0.0015; D1-hM4Di vs D2-hM4Di, p = 0.9999; D1-hM4Di vs D1/D2-mCherry, **p = 0.0041; D2-hM4Di vs D1/D2-mCherry, **p = 0.0036). (F and G) The effects of DREADD suppression on the motivation indices. DREADD suppression of D1-MSN in the NAc decreased the number of earned reward (F) in both the VD-Attend and VD-Avoid task (Left, Two-way RM-ANOVA with Sidak correction, Treatment effects, F_1,12_ = 70.32, ***p < 0.0001; VD-Attend, ***p = 0.0001; VD-Avoid, ***p = 0.0002). DREADD suppression of D2-MSN in the Nac did not affect the number of earned reward in both the VD-Attend and VD-Avoid task (Middle, Two-way RM-ANOVA with Sidak correction, Treatment effects, F_1,14_ = 0.2215, p = 0.6451; VD-Attend, p = 0.7989; VD-Avoid, p = 0.3959). CNO treatment in D1/D2-mCherry did not affect the number of earned reward in both the VD-Attend and VD-Avoid task (Right, Two-way RM-ANOVA with Sidak correction, Treatment effects, F_1,12_ = 0.6391, p = 0.4396; VD-Attend, p = 0.9626; VD-Avoid, p = 0.1759). DREADD suppression of D1-MSN in the Nac increased reward latencies (G) in both the VD-Attend and VD-Avoid task (Left, Two-way RM-ANOVA, Treatment effects, F_1,12_ = 6.259, *p = 0.0278). DREADD suppression of D2-MSN in the Nac did not affect reward latencies in both the VD-Attend and VD-Avoid task (Middle, Two-way RM-ANOVA, Treatment effects, F_1,14_ = 0.8027, p = 0.3854). CNO treatment in D1/D2-mCherry did not affect reward latencies in both the VD-Attend and VD-Avoid task (Right, Two-way RM-ANOVA, Treatment effects, F_1,12_ = 3.705, p = 0.0783).

### Optogenetic Inhibition of D2-MSN in the NAc during Outcome Period on the Error Trial Is Sufficient to Impair the Ability to Execute the Strategy to Avoid a Bad Option on the Next Trial

We have shown that D2-MSN in the NAc causally contributes to the strategy to avoid a bad option (**Figure 2**). To identify a critical time window in which D2-MSN exerts influence on the choice behavior, we optogenetically inhibited the neuronal activity of D2-MSN in the NAc in a time-locked manner. We first bilaterally expressed the light-driven outward proton pump, archaerhodopsin (ArchT) fused to tdTomato (Han et al., 2011) in D2-MSN in the NAc and implanted optic fibers above the NAc (**Figure 3A**, **3B**, **and S4**). The optical stimulation can suppress the activities of ArchT expressing neurons (Kang et al., 2017). We separated the task into 3 time periods (ITI period; the last 5 sec of an inter-trial interval (ITI) to trial initiation. Cue period; the time from trial initiation to the response. Outcome period; 5 sec after the response, **Figure 3C**) to tested whether D2-MSN activity in different time periods have different effects on the performance of the VD-Avoid task. We found that optogenetic inhibition of D2-MSN in the NAc during Outcome period selectively decreased the performance but not in ITI or Cue period stimulation trials (**Figure 3D-3F**). Furthermore, a decrease in performance by optogenetic inhibition during Outcome period was observed on the next trial after making an error choice on the previous trial but not after making a correct choice (**Figure 3G and 3H**). The recent history-dependent decline in the performance was observed only when optogenetic inhibition was applied to Outcome period but not ITI or Cue period (**Figure S5A-S5D**). These results suggest that the neuronal activity of D2-MSN in the NAc immediately after making an error response plays an important role in avoiding a bad option on the next trial. In addition, since response latencies were modulated by recent history (**Figure S1**) and optogenetic inhibition of D2-MSN in the NAc affect the performance in the VD-Avoid task in a history dependent manner (**Figure 3G**), we analyzed whether optogenetic inhibition of D2-MSN in the NAc affected response latencies in the VD-Avoid task. Interestingly, optogenetic inhibition of D2-MSN in the NAc sped up the response on the current trial after having committed an error on the previous trial (**Figure S6A-S6F**), suggesting that the neuronal activation of D2-MSN in the NAc immediately after making an error response contributes to post-error slowing. Taken together, optogenetic inhibition of D2-MSN in the NAc during Outcome period in the error trial selectively decreases the behavioral performance on the next trial.

**Figure 3.**
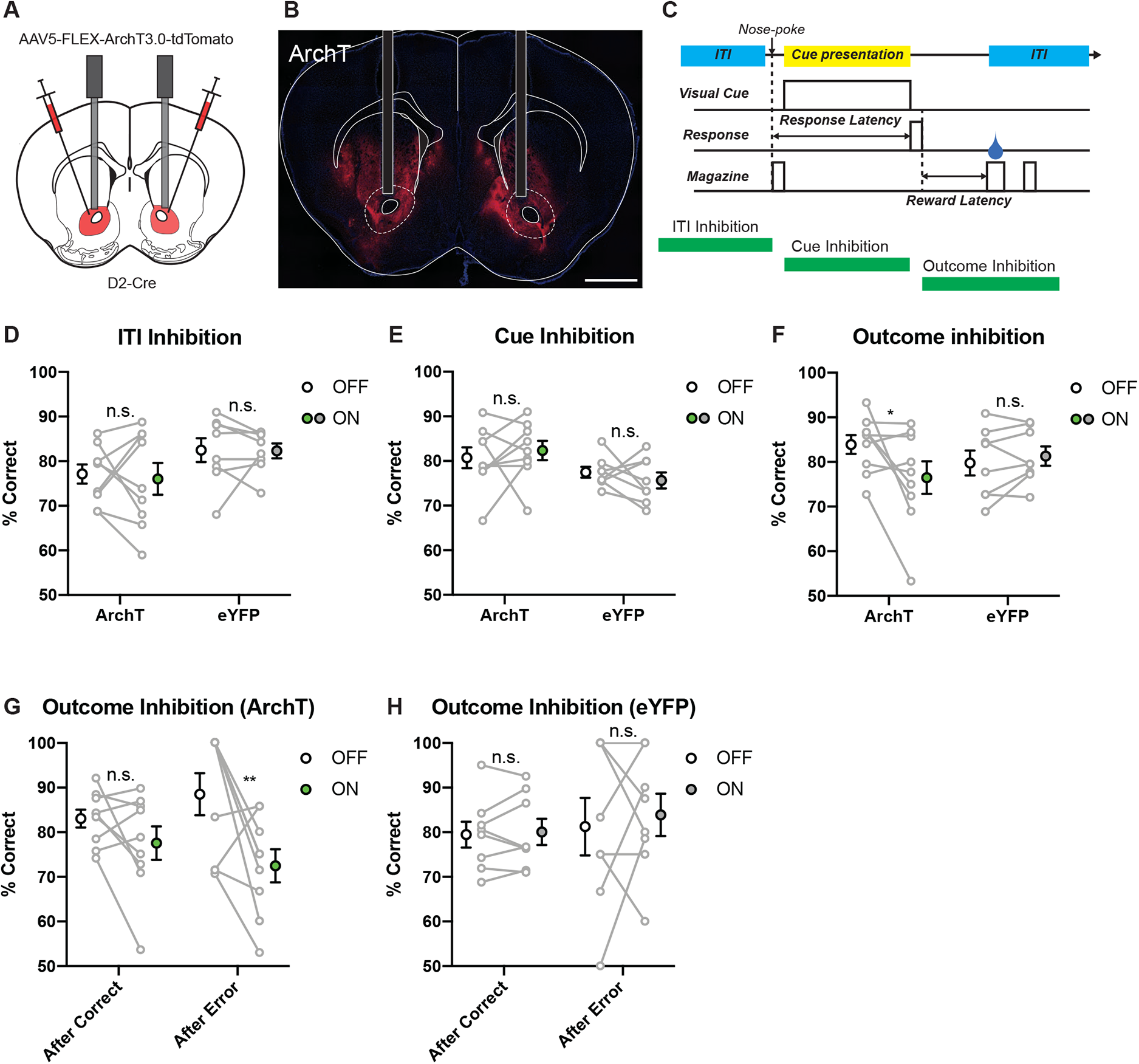
Post-Error Activation of D2-MSN is Necessary for Avoiding a Bad Option. (A) Schematic of viral injection and optic fiber implantation. (B) Representative coronal section with optic fibers. Scale bar, 1 mm. I Schematic of optical stimulation protocol. (D) Optical suppression of D2-MSN in the NAc during ITI period did not affect the behavioral performance (Two-way RM-ANOVA with Sidak correction, Group × Treatment interaction, F_1,15_ = 0.05089, p = 0.8246; ArchT, n = 9, p = 0.9121; eYFP, n = 8, p = 0.9981). (E) Optical suppression of D2-MSN in the NAc during Cue period did not affect the behavioral performance (Two-way RM-ANOVA with Sidak correction, Group × Treatment interaction, F_1,15_ = 0.8700, p = 0.3657; ArchT, p = 0.7815; eYFP, p = 0.7571). (F) Optical suppression of D2-MSN in the NAc during Outcome period decreased the behavioral performance in the next trial (Two-way RM-ANOVA with Sidak correction, Group × Treatment interaction, F_1,15_ = 5.340, *p = 0.0355; ArchT, p = 0.0271; eYFP, p = 0.8368). (G and H) Optical stimulation of D2-MSN in the NAc during Outcome period after error response decreased the behavioral performance of ArchT mice (G) in the next trial (Two-way RM-ANOVA with Sidak correction, Treatment effects, F_1,8_ = 7.134, *p = 0.0283; After Correct, p = 0.2634; After Error, **p = 0.0029) but not eYFP mice (H, Two-way RM-ANOVA with Sidak correction, Treatment effects, F_1,7_ = 0.2682, p = 0.6205; After Correct, p = 0.9909; After Error, p = 0.8477).

### D2-MSN in the NAc Are Dominantly Activated after Making an Error Response than D1-MSN

We have shown that inhibition of D2-MSN in the NAc during Outcome period was sufficient to decrease the behavioral performance in the VD-Avoid task (**Figure 3**). Next, we examined whether D2-MSNs are activated by error response and how the neuronal activity differs between D1-MSN and D2-MSN. To investigate the activity of D1-MSN or D2-MSN in the NAc, we performed calcium imaging of D1-MSN or D2-MSN in the NAc at a single-cell resolution using a miniature microscope. We injected AAV carrying a fluorescent calcium indicator, jGCaMP7f (Dana et al., 2019) in a Cre dependent manner into the NAc of D1-Cre or D2-Cre mice and implanted a gradient-index (GRIN) lens above the NAc (**Figure 4A, 4B, and S7**). Using a constrained non-negative matrix factorization method to the microendoscopic images (CNMFe; Zhou et al., 2018), we extracted the calcium activities from individual D1-MSNs or D2-MSNs from freely moving mice performing the VD-Avoid task (**Figure 4C and 4D; Movies S1 and S2**). We identified 266 cells from D1-Cre mice and 194 cells from D2-Cre mice, respectively. Some of cells were reliably activated after making an error response but not a correct response (**Figure 4E**). Others showed a transient activation following a correct response (**Figure 4F**). We have shown that D1-MSN and D2-MSN contribute to the behavioral performance in a recent history dependent manner (**Figure S3A, S3B, and 4G**). Therefore, we focused on the difference in the calcium activity between correct and error trials and calculated the area under the curve receiver operating characteristic (auROC) of z-scored activities in the error and correct trials. To classify the calcium activities of D1-MSN and D2-MSN in the NAc, we performed the principal component analysis (**Figure 4G**). In D2-MSN, 60% of identified cells (117 out of 194 cells) were classified as Type I, which showed sustained activation after making an error choice while 40% of identified cells (77 out of 194 cells) were classified as Type II, which were activated in a correct trial (**Figure 4H**). In D1-MSN, 44% of identified cells (116 out of 266 cells) were classified as Type I while 56% of identified cells (150 out of 266 cells) were classified as Type II (**Figure 4H**). Moreover, to investigate when D1-MSN or D2-MSN in the NAc showed statistically significant calcium activity between correct and error trials, we defined cell with statistically significant activity as responsive cells and calculated their proportions (**Figure 4I and S8**). As a result, we found that the proportion of responsive cells statistically increased in Outcome period compared to ITI and Cue period (**Figure 4J**). This result was consistent with the previous results by optogenetics in which inhibition of D2-MSN during Outcome period selectively decreased the behavioral performance (**Figure 3D-3F**). Furthermore, we analyzed the calcium dynamics of cells with statistically significant activity during Outcome period (**Figure 4K and S9**) and found that the proportion of Type I (Error Type) in D2-MSN was more dominant than D1-MSN (**Figure 4L**). These results were also consistent with the results by optogenetics in which inhibition of D2-MSN in the error trials decreased the performance on the next trial (**Figure 3G and 3H**). Although Type I and Type II were intermingled in both D1-MSN and D2-MSN (**Figure S10A-S10D**), D2-MSNs of the same valence existed closer to each other than those of opposite valence, suggesting that D2-MSN but not D1-MSN formed spatial clustering of cells with similar functional properties (**Figure S10E and S10F**). Collectively, the majority of D2-MSNs in the NAc was activated by error choice in the VD-Avoid task, corroborating our hypothesis that neuronal activation of D2-MSN in the NAc by error detection plays an important role in avoiding a bad option. On the other hand, the majority of D1-MSNs in the NAc was activated by correct choice. Given that inhibition of D1-MSN in the NAc decreased the performance on the next trial after making a correct choice, it could be that neuronal activation of D1-MSN in the NAc contributes to keep same strategy with confidence.

**Figure 4.**
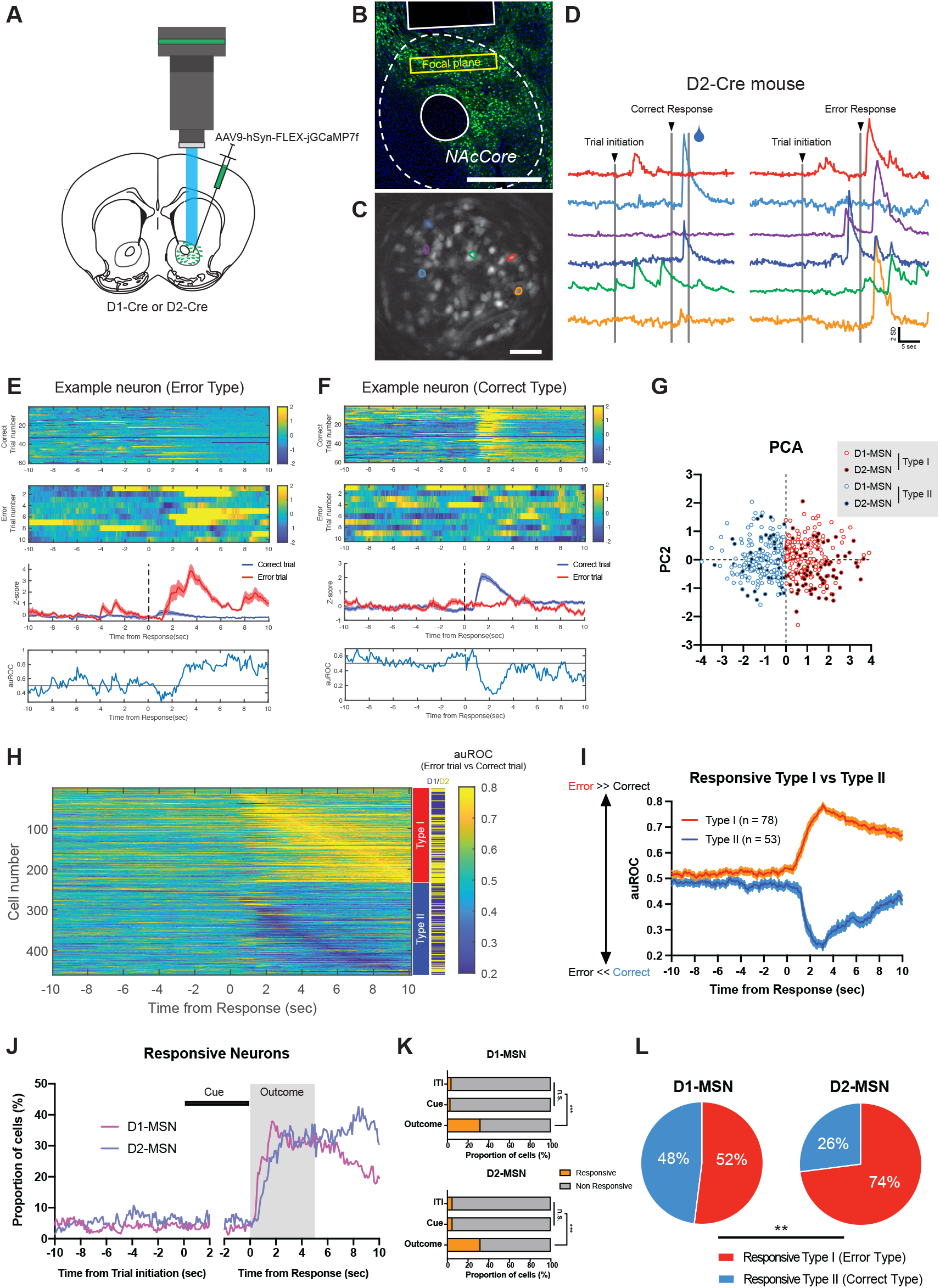
Individual D2-MSNs Are Predominantly Activated by Error Choice. (A) A schematic of viral injection and GRIN lens implantation. (B) Representative coronal image of jGCaMP7f expression in D2-MSNs. Scale bar, 500 µm. (C) Maximum projection image of a representative imaging plane. Scale bar, 50 µm. (D) Example traces of individual neurons from a representative mouse performing the VD-Avoid task. (E) Averaged traces of an example neuron showing error response. Trial-by-trial (Top) and averaged (Middle) responses of an example neuron in correct and error trials. Area under the ROC curve (auROC) reflecting statistical difference between calcium trace of error and correct trials (Bottom). (F) Averaged traces of an example neuron showing correct response. Trial-by-trial (Top) and averaged (Middle) responses of an example neuron in correct and error trials. Area under the ROC (auROC) curves reflecting statistical difference between calcium trace of error and correct trials (Bottom). (G) The first two components of the auROC curves of all the neurons of D1-Cre (n = 3 mice) and D2-Cre mice (n = 3 mice). (H) Heatmaps of the auROC curves of all the neurons of D1-Cre and D2-Cre mice. Each row represents one neuron. (I) Averaged traces of the auROC curves from the responsive Type I neurons (n = 78) or the responsive Type II neurons (n = 53). (J) The proportion of responsive cells across the session. (K) Averaged proportion of responsive cells during ITI, Cue, or Outcome period. The proportion of responsive cells during Outcome period was more than that during ITI or Cue period in both D1-Cre (Top, Chi-square test with Bonferroni correction, χ^2^ = 47.92, ITI vs Cue, p > 0.9999; ITI vs Outcome, ***p < 0.0001; Cue vs Outcome, ***p < 0.0001) and D2-Cre (Bottom, Chi-square test with Bonferroni correction, χ^2^ = 40.37, ITI vs Cue, p > 0.9999; ITI vs Outcome, ***p < 0.0001; Cue vs Outcome, ***p < 0.0001) mice. (L) The fraction of Type I and Type II neurons in D1-Cre or D2-Cre mice. The fraction of Type II neurons in D2-Cre mice were significantly more than that in D1-Cre mice (Chi-square test, χ^2^ = 10.38, **p = 0.0013).

### Neuronal Activation of D2-MSN in the NAc by Error Choice Were Acquired through Learning in the VD-Avoid Task

To investigate if error-correct selective activity during the Outcome period depends on the learning, we next examined whether neuronal activation of MSN in the NAc by error or correct choices in the VD-Avoid task depended on the learning. To investigate how the neuronal activity of D1-MSN or D2-MSN in the NAc changed through learning at a single-cell level, we tracked the same neurons before and after learning (**Figure 5A**) with a cell registration methods (Sheintuch et al., 2017). As the training progressed, D1-Cre and D2-Cre mice similarly improved their performances in the VD-Avoid task (**Figure S11**). In the criterion session, we imaged the activities of 460 cells (D1-Cre, 266 cells; D2-Cre, 194 cells) and identified 239 neuron pairs (D1-Cre, 103 pairs; D2-Cre, 136 pairs) from these mice. A subset of cells activated by error choices (Type I) in the criterion session (expert) had a weak response to error choices in the first session (novice) in the VD-Avoid task (**Figure 5B**). Similarly, we also found that some of the cells activated by correct choices (Type II) in expert mice had a weak response to correct choices in novice mice in the VD-Avoid task (**Figure 5C**). Next, to investigate the change in the calcium activity against error or correct choices through learning, we analyzed the difference in the calcium activities in novice and expert mice (**Figure 5D**). By tracking the same neurons through training, we found that the activity of D2-MSN in expert mice positively correlated with that in novice mice, whereas we could not detect any correlation in D1-MSN imaged in the VD-Avoid task (**Figure 5D**). Furthermore, we classified the type of neuron pairs and quantified the proportion of cells that become responsive Type I or Type II from non-responsive in D1-Cre or D2-Cre mice. The results showed that the majority (13.6%) of D1-MSN acquired the correct choice selectivity after learning (**Figure 5E and S12**). In contrast, the majority (16.8%) of D2-MSN acquired the error choice selectivity after learning (**Figure 5E and S12**). Moreover, we revealed that the proportion of cells that become responsive Type I from non-responsive were significantly higher in D2-MSN than D1-MSN (**Figure 5F**). In contrast, the proportion of cells that become responsive Type II from non-responsive were significantly higher in D1-MSN than D2-MSN (**Figure 5F**). These results indicate that the neuronal activation of D1-MSN and D2-MSN by outcome depends on the learning. The acquisition of responsiveness by learning of these neurons is likely to contribute to the high performance in the VD-Avoid task.

**Figure 5.**
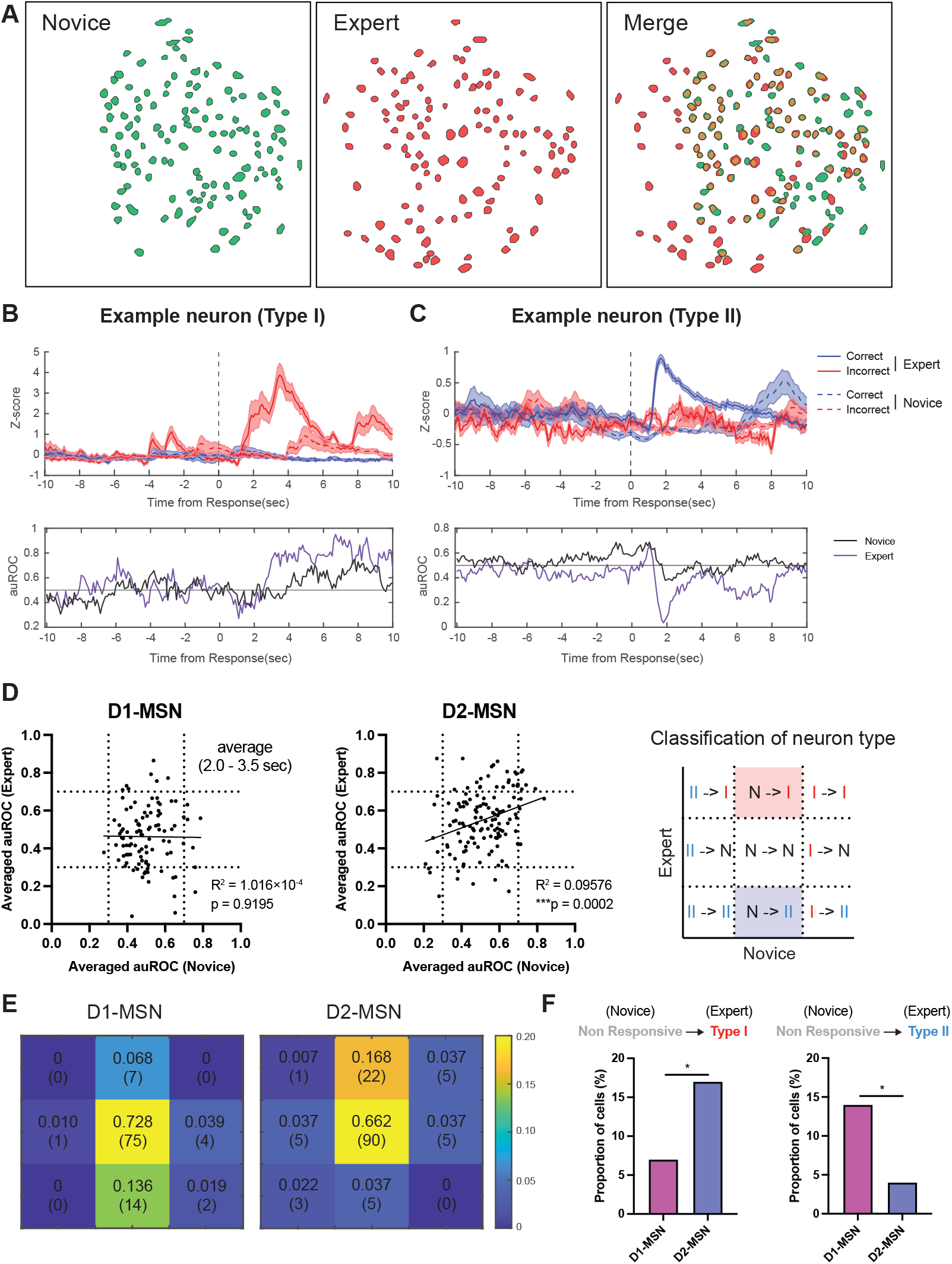
More Non-Responsive D2-MSNs Acquire Error Responses than D1-MSNs through Learning. (A) Representative image of contour map from novice (green) and expert (red). (B) Averaged traces of an example neuron acquiring error response through learning. Averaged calcium traces of novices and experts in correct and error trials (Top). Area under the ROC curve (auROC) reflecting statistical difference between calcium trace of error and correct trials from novices and experts (Bottom). (C) Averaged traces of an example neuron acquiring correct response through learning. Averaged calcium traces of novices and experts in correct and error trials (Top). Area under the ROC curve (auROC) reflecting statistical difference between calcium trace of error and correct trials from novices and experts (Bottom). (D) Relationship between averaged auROC of novice and that of expert in D1-Cre (Left, n = 3 mice) or D2-Cre mice (Middle, n = 3 mice). Schematic of classification of neuron type (Right). I, Type I; II, Type II; N, Non-responsive. (E) The fraction of each neuron type in D1-Cre (Left) and D2-Cre mice (Right). (F) Non-responsive D2-MSN preferentially became Type I (Error Type) through learning (Left, Chi-square test, χ^2^ = 4.735, *p = 0.0296), while Non-responsive D1-MSN preferentially became Type II (Correct Type) through learning (Right, Chi-square test, χ^2^ = 6.105, *p = 0.0135).

## DISCUSSION

Although dopamine signaling plays an important role in strategy-dependent decision making (Frank et al., 2004, 2007), it has been poorly understood how the neural substrates targeted by dopamine contribute to reinforcement learning and execution of the strategy under the decision. Here we have established novel visual discrimination tasks in mice to assess the neural mechanisms underlying the strategy-dependent decision making. Our data show that neuronal activation of D2-MSN in the NAc by error choices is essential for executing the strategy to avoid a bad option. Previous studies have shown that when the dopamine concentration in a brain is increased by L-dopa treatment, the subjects cannot avoid a bad option but still can choose a good choice (Frank et al., 2004, 2007). L-dopa treatment can decrease the excitability of D2-MSN because the dopamine D2 receptor is coupled to Gi signaling (Cepeda et al., 1993; Stoof and Kebabian, 1984; Surmeier et al., 2007). Given that our chemogenetic and optogenetic suppression of D2-MSN in the NAc disrupted the strategy to avoid a bad option but not that to choose a good option, we concluded that hypoactivity of D2-MSN in the NAc caused the subjects not to execute the strategy to avoid a bad option. Although overactivation of D1-MSN in the striatum including the NAc could also contribute to disability to avoid a bad option directly or indirectly (e.g. via local D2-MSN to D1-MSN synapse; Matamales et al., 2020), our study demonstrates downstream neural mechanisms of dopamine signaling by which the neuronal activity of D2-MSN in the NAc selectively contributes to avoid a bad option but not choose a good option.

Our calcium imaging data suggest that as individual D2-MSNs are reliably activated by error choices, mice efficiently avoid a bad option. Although individual D1-MSNs lost the response selectivity through learning (**Figure 5D**), given that chemogenetic suppression of D1-MSN decreased the performance in the VD-Avoid task as well as the VD-Attend task (**Figure 2D**) and some D1-MSNs were recruited by correct choices after learning (**Figure 5E and 5F**), we expect that the neuronal activity of D1-MSN also contributes to the performance in the VD-Avoid task. Our calcium imaging data also showed that the activity of D2-MSN in expert mice positively correlated with that in novice mice, whereas we could not detect any correlation in D1-MSN imaged in the VD-Avoid task. Given that the VD-Avoid task required mice to avoid a specific cue (e.g. flag) associated with negative outcome (e.g. omission), the neuronal activation of D2-MSN in the NAc by error choices plays an important role in avoiding a bad option, and the neuronal activity of D2-MSN had a positive correlation between before and after learning, individual D2-MSNs in the NAc could encode the value information associated with the specific cue, in particular negative value. Moreover, given that D1-MSN plays an important role in a reward learning and a reward approaching behavior (Hikida et al., 2010; Kravitz et al., 2012; Macpherson and Hikida, 2018; Shin et al., 2018; Tai et al., 2012), the neuronal activation of D1-MSN by positive outcome (e.g. reward) can contribute to choose the cue associated with positive outcome. The reason we could not find any correlation in D1-MSN was attributed to the property of the VD-Avoid in which mice could not associate the specific cue with the positive outcome because the cue associated with reward changed in every trial in the VD-Avoid task.

Previous studies indicate that D2-MSN plays an important role in aversive learning (Danjo et al., 2014; Hikida et al., 2010). However, classical paradigms trained mice not to take any action against the cue or the context associated with an aversive stimulus. Therefore, it was difficult to distinguish the motor component from an aversive information processing because D2-MSN is also involved in action and locomotion (Bakhurin et al., 2020; Kravitz et al., 2010; Tecuapetla et al., 2016; Tsutsui-Kimura et al., 2017a; Yao et al., 2021). Furthermore, although an electric shock or an air puff was often used as an aversive stimulus, those stimuli are innately aversive and could be different from a negative prediction error (e.g. omission). Because our paradigm shares the other components from the strategy, we think that it is easier to extract the component of the strategy. Our previous data showed that perturbation of the D2 receptor signalling by knockout of D2L interfered with visual discrimination learning (Morita et al., 2016). By combining the previous study and the current study, we suggest that mice simultaneously learn to choose a cue associated with a reward and to avoid a cue not associated with a reward in classical discrimination learning.

It is known that dopamine neurons encode a reward prediction error and the firing rate of dopamine neurons decreases (e.g. dopamine dip) when the subjects did not receive the expected reward (Schultz et al., 1997). We expect that the dopamine dip by a reward omission contributes to the strategy to avoid a bad option. Activation of D2-MSN by disinhibition of D2 receptor could contribute to suppressing the action associated with a negative outcome under discrimination learning. Interestingly, dopamine neurons are heterogeneous, and reward prediction error activity can be observed in the NAcCore but not NAcShell (Iino et al., 2020; Jong et al., 2019). We mainly targeted the NAcCore and revealed that chemogenetic and optogenetic suppression of D2-MSN in the NAcCore is sufficient to decrease the performance in the VD-Avoid task. Although we cannot exclude the possibility that the NAcShell also contributes to the strategy to avoid a bad option, we revealed that at least D2-MSN in the NAcCore plays an important role in the strategy to avoid a bad option. Moreover, it is reported that the dorsomedial striatum (DMS) plays an important role in reinforcement learning (Kravitz et al., 2012; Tai et al., 2012) and electrophysiological recording showed that D1-MSN in the DMS preferentially encodes positive prediction error and D2-MSN encodes negative prediction error (Shin et al., 2018). Therefore, the DMS could also contribute to the strategy. In the future, we need to investigate how differently the NAcCore, NAcShell, and DMS contribute to the execution of the strategy under decision.

Disability to execute the strategy to avoid a bad option could be associated with risk-taking behaviors. A previous study showed D2-MSN activity in the NAc is important for avoiding risk-taking behavior (Zalocusky et al., 2016). They showed D2-MSN activation in the NAc contributes to avoiding a risky choice. Our results basically support their data and extend it to that D2-MSN activity in the NAc is involved in suppressing the action associated with the negative outcome as well as a risky choice. However, they showed D2-MSN activation occurs before the decision in the trial after the failure of risky-choice and optogenetic activation of D2-MSN during the decision period is sufficient to avoid a risky choice. Our data showed that optogenetic inhibition of D2-MSN in the NAc during ITI or Cue period did not affect the decision. Although the exact mechanism is unknown because they did not test the effect of optogenetic activation of D2-MSN in the NAc during outcome period on a risky choice, the timing of activation of D2-MSN in the NAc may differ depending on the task. Moreover, their target is a medial region of the NAc, while our target is the lateral region. Because dopamine dynamics are different depending on the region of the NAc (Iino et al., 2020; Jong et al., 2019), the difference in activation timing may depend on the region. However, further investigations are needed in the future.

Interestingly, our results showed that optogenetic inhibition of D2-MSN in the NAc during ITI or Cue period did not affect the performance, suggesting that D2-MSN activity does not contribute to discrimination itself. Those results also suggest that the action value information is transmitted from the NAc via the thalamocortical loop and encoded in the neurons in different areas such as the parietal cortex (Akrami et al., 2018) and the DMS (Tai et al., 2012) during the decision period. Moreover, because it is known that D2-MSN in the NAc sends inhibitory input onto the VP (Gallo et al., 2018) and inhibition of GABAergic neurons in the VP reduces the risky choice (Farrell et al., 2021), inhibition of GABAergic neurons in the VP via D2-MSN in the NAc could be important for avoiding a bad option.

A previous study has shown that inhibition of D2-MSN in the NAc could lead to rewarding effects (Gallo et al., 2018). However, our results cannot be explained by simple rewarding effects because optogenetic inhibition of D2-MSN in the NAc selectively sped up the response latency in the trial after error. If the rewarding effect by optogenetic inhibition affects the performance in the VD-Avoid task, the response latency in the trial after correct should be similarly decreased. Given that activation of D2-MSN in the NAc increases the motivation, D2-MSN in the NAc could be involved in decision making via complex mechanisms. This history-dependent effect on response latencies is interesting from a psychological point of view. A number of psychological studies has reported that subjects tend to slow down the response on the current trial after having committed an error on the previous trial (Houtman et al., 2012; Notebaert et al., 2009; Rabbitt and Rodgers, 1977; Ruitenberg et al., 2014). Although the neural mechanisms underlying the post-error slowing have been not fully understood, in this study we have shown that inhibition of D2-MSN in the NAc sped up the response latency in the trials after error choice. The result suggests that activation of D2-MSN in the NAc after error choice contributes to the post-error slowing.

Contradictory to a classical model, our results showed a subset of D1-MSNs in the NAc encodes the negative information. However, given that it is reported that a subset of D1-MSN in the dorsal striatum selectively encodes the aversive information (Kim et al., 2020; Xiao et al., 2020), the NAcCore could also include the similar subset of D1-MSN. Given that majority of D1-MSN in the NAc were activated by reward and bulk inhibition of D1-MSN decreased the motivation in our experiments, we conclude that D1-MSN in the NAc mainly plays an important role in motivation. However, interestingly, chemogenetic suppression of D1-MSN in the NAc affected the performance in a recent history-dependent manner. This result cannot be explained by the simple effect of decreased motivation. Although further experiments are required, we think that there are at least three subpopulations in D1-MSN of the NAc. The first population is involved in a general motivation control. The second modulates the behavior based on reward history. The third encodes a negative prediction error. Although we could not analyze the contribution of D1-MSN to the strategy because bulk suppression of D1-MSN in the NAc drastically decreased the motivation, given that D1-MSN in the NAc plays an important role in approaching behaviors in the Pavlovian conditioning (Macpherson and Hikida, 2018), we expect that a subset of D1-MSN in the NAc contribute to the strategy to choose a good option. Considering that D2-MSN in the NAc were activated by error and inhibition of D2-MSN in the NAc decreased the performance in the trials after error response, D1-MSN and D2-MSN in the NAc have opposite functions in the visual discrimination learning and activation of D1-MSN by reward and activation of D2-MSN by error could lead to making the same choice (e.g. via high confidence) and avoid the previous choice, respectively. Moreover, D2-MSN is involved in reversal learning and switching behaviors (Macpherson et al., 2016; Matamales et al., 2020; Tsutsui-Kimura et al., 2017b; Yawata et al., 2012). Because these behaviors are modulated by negative prediction error (e.g. reward omission), the strategy to avoid a bad option is required. Although further experiments are needed to test if the same population regulate the strategy to avoid a bad option and reversal learning, we conclude that D2-MSN in the NAc is important for detecting a negative outcome and avoiding the action associated with the negative outcome.

It is known that decision strategy could be influenced by a brain state and maladaptive decision making can be observed in various neurological and psychiatric disorders such as Parkinson’s disease, drug addiction, and mood disorders (Frank et al., 2004; Hikida et al., 2010; Kunisato et al., 2012; Strickland et al., 2016). In Parkinson’s disease, it is known that cell death of dopamine neurons occurs and acute dopamine depletion leads to overactivation of D2-MSN (Parker et al., 2018). Moreover, a study shows that repeated cocaine treatment reduced the frequency of miniature excitatory postsynaptic currents in D2-MSN (Kim et al., 2011). These studies suggest that increase in excitability of D2-MSN leads to the strategy to avoid a bad option, while decrease in excitability of D2-MSN causes to disability to avoid a bad option. These bidirectional effects on the strategy support our hypothesis that activation of D2-MSN plays an important role in avoiding a bad option. Although it is difficult to explain with one factor because the plasticity change occurs at neural circuit level in various neurological and psychiatric disorders (Macpherson and Hikida, 2019), it is likely that excitability of D2-MSN contributes to the strategy to avoid a bad option. In contrast to L-dopa treatment, depression selectively disrupts the strategy to choose a good option (Kunisato et al., 2012). Because depression is strongly related to the concentration of serotonin in the brain (Warden et al., 2012), the abnormal neural activity of a subset of the neuron in the NAc via serotonin receptor could be associated with the disability to execute the strategy to choose a good option.

In conclusion, we provide the evidence in which D2-MSN in the NAc selectively contributes to the strategy to avoid a bad option. Moreover, activation of D2-MSN in the NAc by error choices plays an important role in acquisition of the strategy to avoid a bad option. Understanding the neural mechanisms underlying the strategy will help to treat with dysfunctions to execute the appropriate strategy in various psychiatric disorders such as Parkinson’s disease, drug addiction, and mood disorders. In addition, the development of VD-Attend and VD-Avoid task could lead to a useful novel model to understand the neural mechanisms by which different people execute different strategies depending on the genes, situations, developments, and ages.

## ACKNOWLEDGEMENTS

This work was supported by MEXT/JSPS KAKENHI grants numbers, JP21K15209 (to T.N.), JP21K15210 (to T.M.), JP19H04983 and JP21H02804 (to K.H.), JP16H06568 and JP18H02542 (to T.H.), by AMED under grant number JP21wm0425010 (to T.H.), by Takeda Life Science Research Foundation (to T.H.), by the Salt Science Research Foundation (No. 2137 to T.H.), by Taiju Life Social Welfare Foundation (to T.H.), and by Smoking Research Foundation (to T.H.). We thank members of Hikida laboratory for maintenance of animals and helpful discussion.

## AUTHOR CONTRIBUTIONS

T.N. conceived the project, conducted all the experiments, performed the analysis, and wrote the manuscript. T.M. supported calcium imaging experiments. K.H. and T.H. contributed to the analysis. T.H. supervised the project. T.M., K.H., and T.H. reviewed and edited the manuscript.

## DECLARATION OF INTERESTS

The authors declare no competing financial interests.

## METHOD DETAILS

### Animals

Wild-type C57BL/6J mice (male, 8-10 weeks old) were used for validation of behavioral experiments. For DREADD, optogenetic, and calcium imaging experiments, male heterozygous D1-Cre (FK150Gsat) and D2-Cre (ER44Gsat) mice were used (DREADD, D1-Cre, n = 12, one mouse was excluded due to no viral expression; D2-Cre, n = 12, one mouse was excluded due to unstable behavior; optogenetics, D2-Cre, n = 19, 2 mice were excluded due to insufficient conditioning; calcium imaging, D1-Cre, n = 3; D2-Cre, n = 4, one mouse was excluded because GRIN lens was located anterior to the NAc). D1-Cre and D2-Cre were maintained in a C57BL/6J background. Animals were housed on a 12-hour light/dark cycle. Behavioral studies were conducted during the light cycle. Mice were kept on water restriction during behavioral testing. For all behavioral experiments except for calcium imaging experiment, mice were grouped housed throughout the experiments. For calcium imaging experiment, mice were singly housed after GRIN lens implantation. All experiments conformed to the guidelines of the National Institutes of Health experimental procedures, and were approved by the Animal Experimental Committee of Institute for Protein Research at Osaka University (approval ID 29-02-1).

### Stereotaxic Surgery

All mice used in this study were anesthetized with ketamine (100 mg/kg) and xylazine (20 mg/kg) for surgical procedures and placed in a stereotaxic frame (Kopf Instruments).

For DREADD experiments, heterozygous D1-Cre or D2-Cre mice were bilaterally injected with 400 nl of AAV5-hSyn-DIO-hM4Di(Gi)-mCherry (5.1×10^12^ GC/ml, UNC) or AAV5-hSyn-DIO-mCherry (5.2×10^12^ GC/ml, UNC) using a Nanoject II instrument (Drummond) at a rate of 100 nl/min (coordinates in mm: AP +1.20, ML ±1.25 from bregma, and DV −3.50 from brain surface. The injection pipette remained in place for 5–10 min to reduce backflow.

For optogenetics experiments, heterozygous D2-Cre mice were bilaterally injected with 400 nl of AAV5-CAG-FLEX-ArchT3.0-tdTomato (1.3×10^13^ GC/ml, Addgene) or AAV5-EF1a-DIO-eYFP (1.3×10^13^ GC/ml, Addgene) were using a Nanoject III instrument (Drummond) at a rate of 100 nl/min (coordinates in mm: AP +1.20, ML ±1.25 from bregma, and DV −3.50 from brain surface. The injection pipette remained in place for 5–10 min to reduce backflow. After retraction, 200 µm diameter (NA 0.37) optic fibers (Thorlabs) were bilaterally implanted with the dental cement (Superbond) at AP +1.20, ML ±1.30 from bregma, and DV −3.20 from brain surface.

For calcium imaging experiments, heterozygous D1-Cre or D2-Cre mice were unilaterally injected with 1200 nl of AAV9-FLEX-jGCaMP7f (9.6×10^12^ GC/ml, Addgene) were stereotaxically injected using a Nanoject III instrument (Drummond) at a rate of 100 nl/min (coordinates in mm: AP +1.20, ML ±1.25 from bregma, and DV −3.60 and −3.10 from brain surface. The injection pipette remained in place for 5–10 min to reduce backflow. After virus injection, a sterile 21-gauge needle was slowly lowered into the brain to a depth of −2.0 mm from the brain surface to aspirate brain tissue above the NAc. The GRIN lens (600 µm diameter, Inscopix) was slowly lowered into the brain to a depth of −3.20 mm from the brain surface by using a GRIN lens holder (Inscopix). We secured the GRIN lens to the skull with the dental cement (Superbond). A silicone elastomer (Kwik-Cast; World Precision Instruments) was applied to the top of the lens to prevent an external damage. Four to six weeks after lens implantation, a baseplate (Inscopix) attached to the miniature microscope (nVista; Inscopix) was positioned above the GRIN lens. The focal plane was adjusted on the basis of the position that blood vessels were clearly observed. After positioning adjustment, the baseplate was secured with the dental cement.

### Behavioral Experiments

#### Apparatus

Training and testing were conducted in a Bussey-Saksida touchscreen chamber (Lafayette Instrument). A black plastic mask with 2 windows (70×75 mm^2^ spaced, 5 mm apart, 16 mm above the floor) was placed in front of the touchscreen. ABET II and WhiskerServer software (Lafayette) were used to control operant system and data collection.

#### Pretraining

As the first phase (3 days), mice were habituated to the chamber in 40-min sessions. Diluted condensed milk (7 µl, Morinaga Milk) was dispensed in the food magazine every 10 sec. In the following phase (1 day), a stimulus was randomly displayed in one of the 2 windows. After a 30-sec stimulus presentation, the milk reward (20 µl) was delivered with a tone (3 kHz) and magazine light. When mice collected the reward, the magazine light went out, and the next trial commenced (60 trials, or up to 60 min) with a new stimulus after a 20-sec intertrial interval (ITI). In the next phase, stimuli were randomly displayed in one of 2 windows, and mice were obligated to touch the stimulus to receive a reward. In the final phase of the pretraining, when a blank window was touched, mice were punished with a 5-sec time-out. After reaching criterion (77% correct for 2 consecutive days), mice moved on to basic training.

#### Basic training

Mice were tested 5–6 days per week (60 trials per day, or up to 60 min). Each trial was initiated after mice nose-poked in the magazine. The visual cues were presented until mice respond to any windows.

For VD-Attend task, two visual cues (Marble and a random image) were presented in the touchscreen. A random image was pseudorandomly chosen from 51 images. If a mouse responded to visual cue “Marble”, milk reward (7 µl) was delivered with a tone (3 kHz) and magazine light. When mice collected the reward, the magazine light went out, and the next trial commenced (60 trials, or up to 60 min) with a new stimulus after a 20-sec intertrial interval (ITI). If a mouse responded to visual cue “a random image”, the mouse was punished with a 5-sec time-out (house light on).

For VD-avoid task, two visual cues (Flag and a random image) were presented in the touchscreen. If a mouse responded to visual cue “a random image”, milk reward (7 µl) was delivered with a tone (3 kHz) and magazine light. If a mouse responded to visual cue “Flag”, the mouse was punished with a 5-sec time-out (house light on).

A response to a random image was recorded as a correct response, while a response to visual cue “Flag” was recorded as an incorrect response.

After reaching criterion (>80% correct for 2 consecutive days), mice moved on to probe test for DREADD experiments, cable habituation for optogenetic experiments, respectively.

For DREADD experiments, vehicle at day1, 2, 3, 5, or CNO (1.0 mg/kg diluted with vehicle, Sigma Aldrich) at day4, 6 was intraperitoneally administered 30 min before the session.

For optogenetic inhibition experiments, once the performance stabilized (>80% correct for 2 consecutive days) with cable, optogenetic stimulation starts. LED power was set to 1-3 mW. Stimulation schedule was counterbalanced.

For calcium imaging experiments, data was acquired at 20 Hz with 0.6 mW LED at the first session of the basic training (Novice) and the session after reaching criterion (the criterion session; Expert). After acquisition, calcium recording files were temporally (factor of 2) and spatially (factor of 4) downsampled and motion-corrected using Inscopix Data Processing software ver 1.3.0. The fluorescent traces of individual neurons were extracted from these images by CNMFe (Zhou et al., 2018). Z-scores were calculated from all recording data.

Percentage correct (correct trials divided by correct plus incorrect trials, recorded as percent), and latencies to correct response, incorrect response, and reward collection were monitored in all behavioral experiments.

### Histology

Animals were deeply anesthetized and transcardially perfused with 0.01 M PBS followed by 4% paraformaldehyde (PFA) in 0.1 M PB (pH 7.4). Brains were removed and post-fixed with 4% PFA at 4 °C for 2 days. After cryoprotection, brains were embedded in OCT compound and cryosectioned (40 µm). Sections were mounted with antifade mouting medium with DAPI (Vectashield). Stitched images were acquired using a Keyence BZ-X800 microscope.

### Statistical analyses

Prism (Graphpad) software was used for statistical analyses. The behavioral performances in wild-type were analyzed using unpaired t-test. DREADD data were analyzed using two-way RM ANOVA with Group (hM4Di, mCherry) and Drug Treatment (vehicle, CNO) or one-way ANOVA (D1-hM4Di, D2-hM4Di, D1/D2-mCherry). Optogenetic data were analyzed using two-way RM ANOVA with Group (ArchT, eYFP) and Light stimulation (OFF, ON) or History (After Correct, After Error) and Light stimulation (OFF, ON). Post hoc Sidak’s multiple comparisons test was performed when F-ratios were significant (p < 0.05). Comparisons of proportion of cells were made using chi-squared test. Simple linear regression was made to calculate the correlation of auROC between novice and expert. Frequency distributions were compared using the Kolmogorov-Smirnov (KS) test. All data are expressed as means ± SEM.

---

**Supplementary Figure S1.**
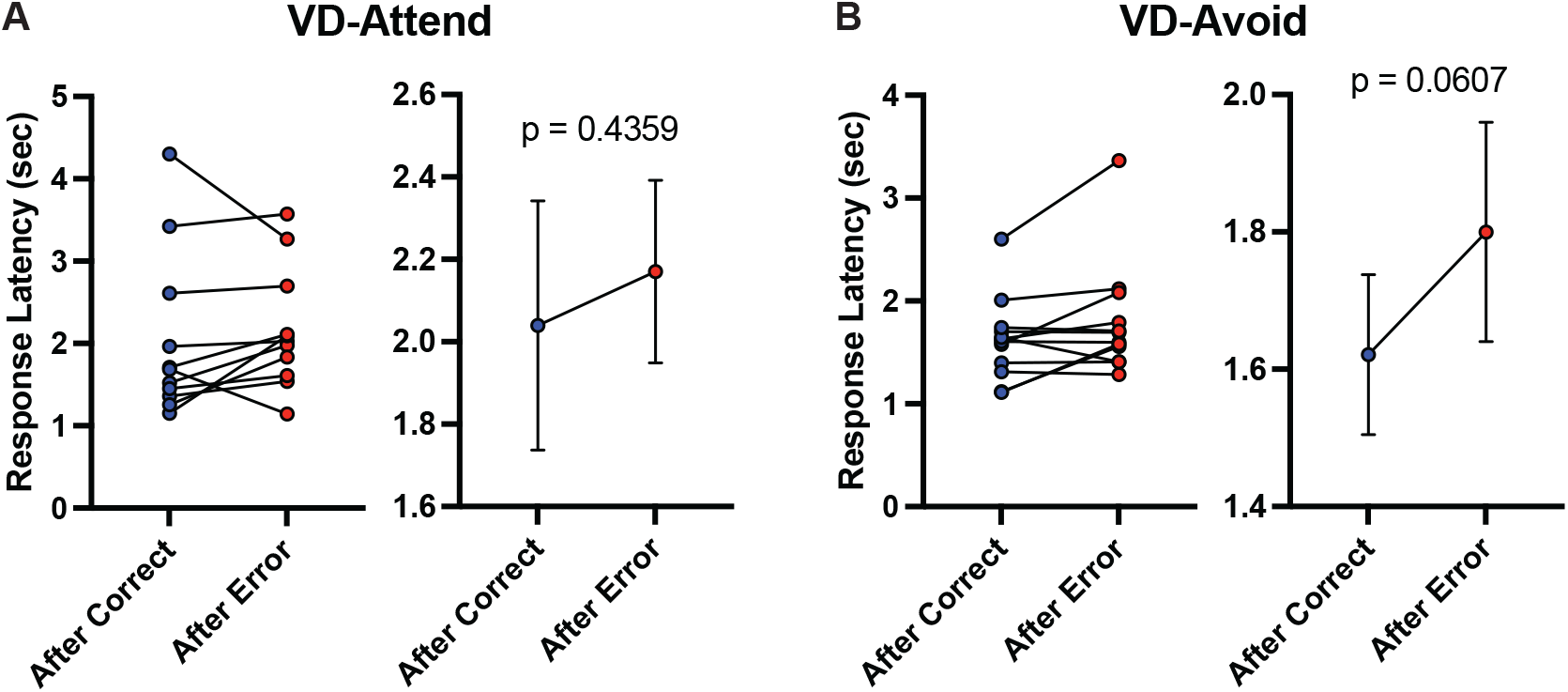
Response Latencies Were Modulated by Recent History. (A) The effects of recent history on response latencies in the VD-Attend task (unpaired t-test, t_10_ = 0.8115, p = 0.4359). (B) The effects of recent history on response latencies in the VD-Avoid task (unpaired t-test, t_11_ = 2.090, p = 0.0607).

**Supplementary Figure S2.**
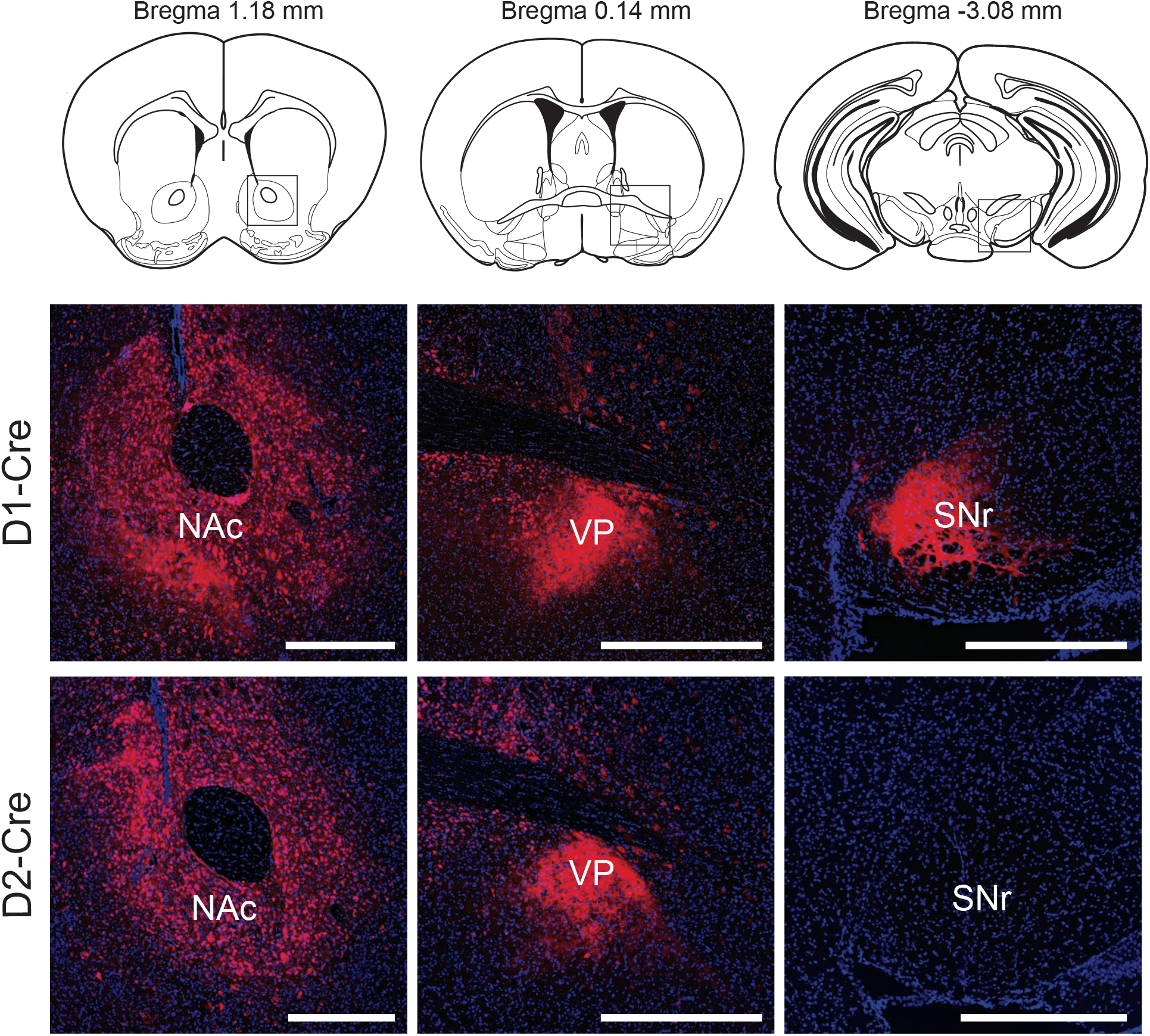
Cell Type Specific Gene Expression. Representative coronal sections of DREADD expression in the D1-MSN or D2-MSN of the NAc (Left). Representative coronal sections containing projection areas, ventral pallidum (VP, Middle) and substantia nigra pars reticulata (SNr, Right).

**Supplementary Figure S3.**
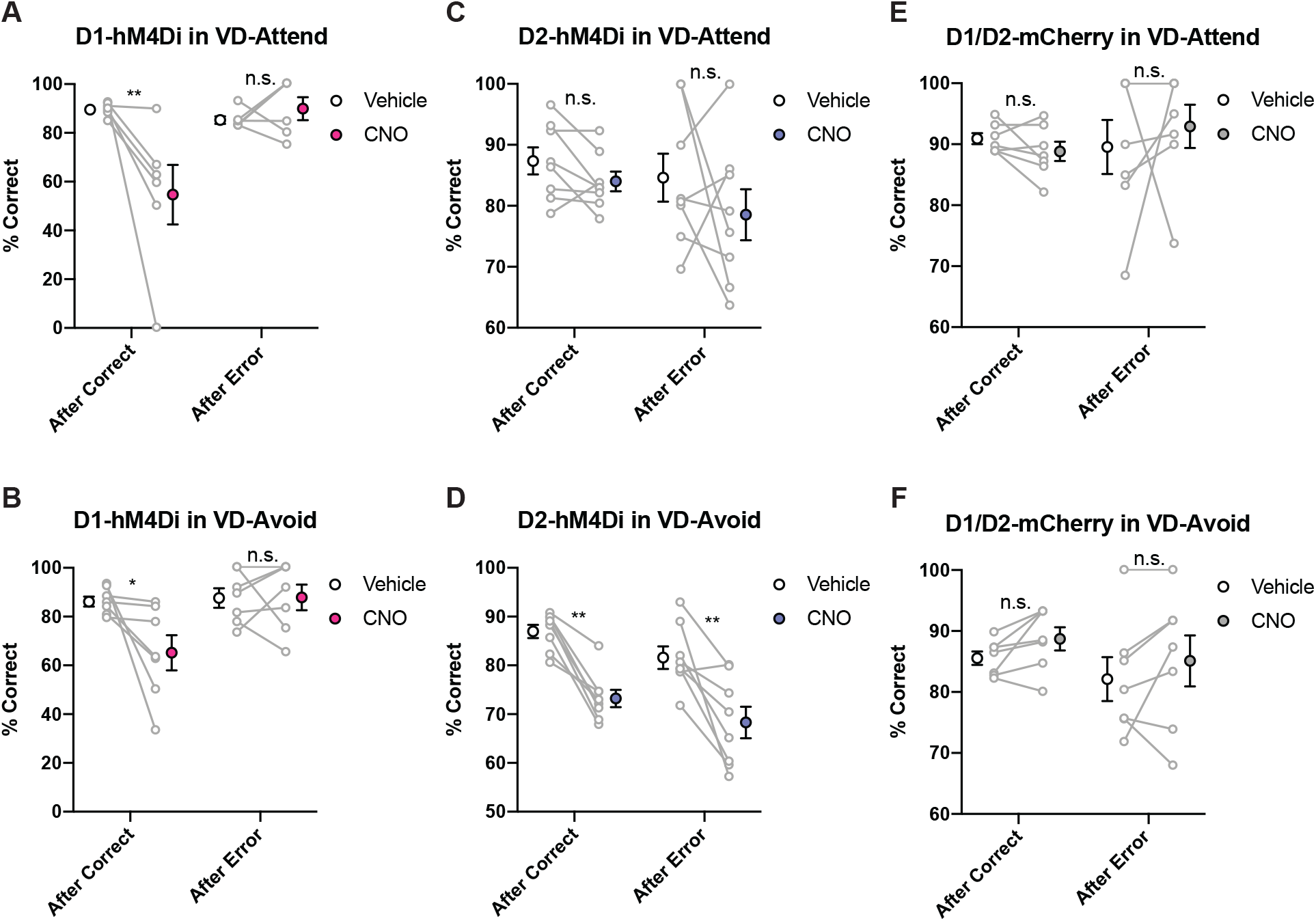
DREADD Suppression of D1-MSN in the NAc Decreases the Behavioral Performance in a Recent History Dependent Manner. (A) DREADD suppression of D1-MSN in the NAc decreased the behavioral performance after correct trial in the VD-Attend task (Two-way RM-ANOVA with Sidak correction, Trial type × Treatment interaction, F_1,11_ = 8.323, *p = 0.0148; After Correct, **p = 0.0015; After Error, p = 0.6357). (B) DREADD suppression of D1-MSN in the NAc decreased the behavioral performance after correct trial in the VD-Avoid task (Two-way RM-ANOVA with Sidak correction, Trial type × Treatment interaction, F_1,12_ = 5.603, *p = 0.0356; After Correct, *p = 0.0133; After Error, p = 0.9990). (C) DREADD suppression of D2-MSN in the NAc did not affect the behavioral performance after correct or error trial in the VD-Attend task (Two-way RM-ANOVA with Sidak correction, Trial type × Treatment interaction, F_1,14_ = 0.3984, p = 0.5381; After Correct, p = 0.7098; After Error, p = 0.3406). (D) DREADD suppression of D2-MSN in the NAc decreased the behavioral performance after correct and error trial in the VD-Avoid task (Two-way RM-ANOVA with Sidak correction, Treatment effects, F_1,14_ = 29.79, ***p < 0.0001; After Correct, **p = 0.0012; After Error, **p = 0.0014). (E) CNO treatment in D1/D2-mCherry did not affect the behavioral performance after correct or error trial in the VD-Attend task (Two-way RM-ANOVA with Sidak correction, Trial type × Treatment interaction, F_1,12_ = 1.058, p = 0.3239; After Correct, p = 0.8609; After Error, p = 0.6787). (F) CNO treatment in D1/D2-mCherry did not affect the behavioral performance after correct or error trial in the VD-Avoid task (Two-way RM-ANOVA with Sidak correction, Trial type × Treatment interaction, F_1,12_ = 0.001573, p = 0.9690; After Correct, p = 0.7057; After Error, p = 0.7319).

**Supplementary Figure S4.**
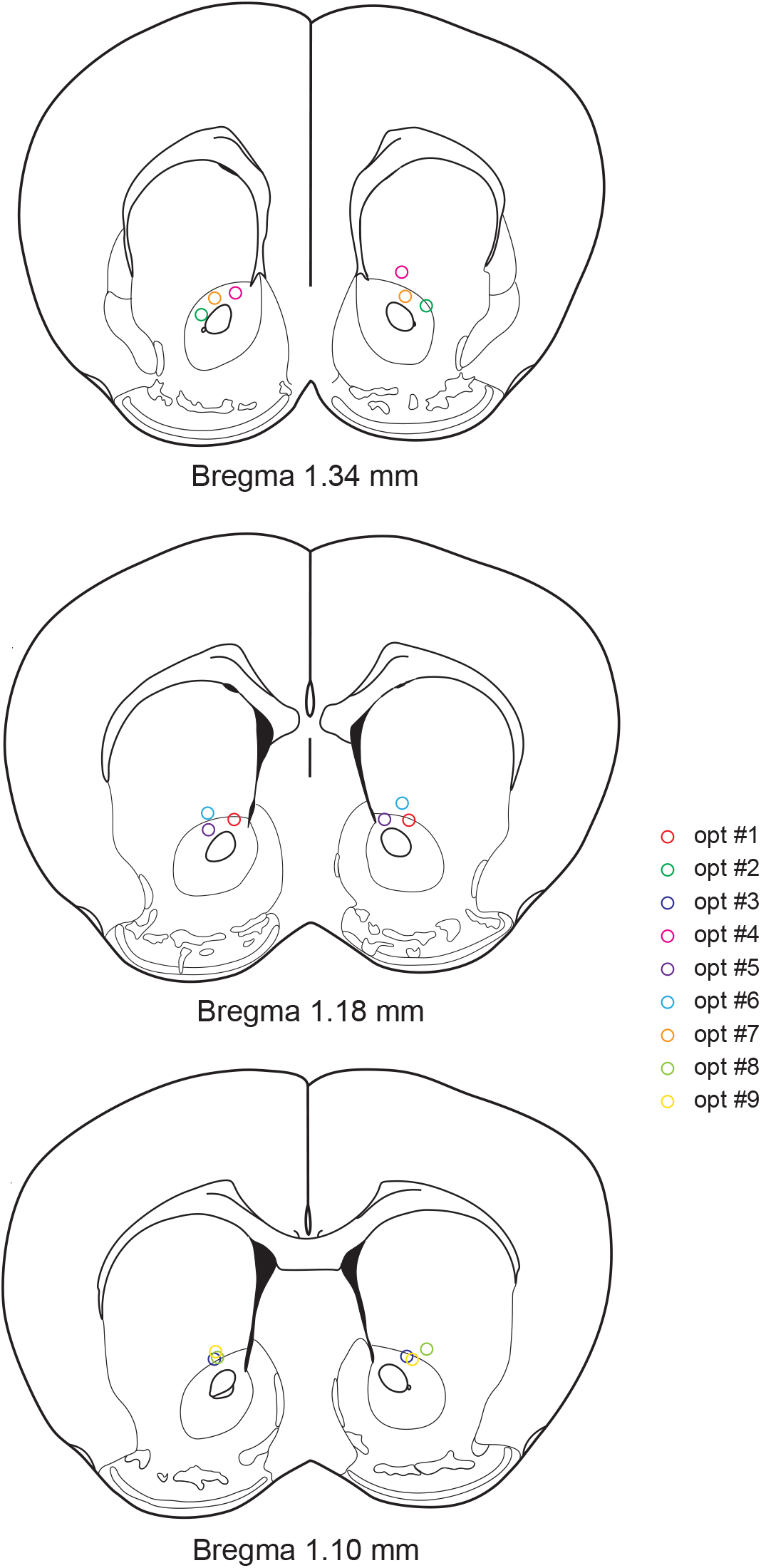
Optic Fiber Placement. Histology on optic fiber placement.

**Supplementary Figure S5.**
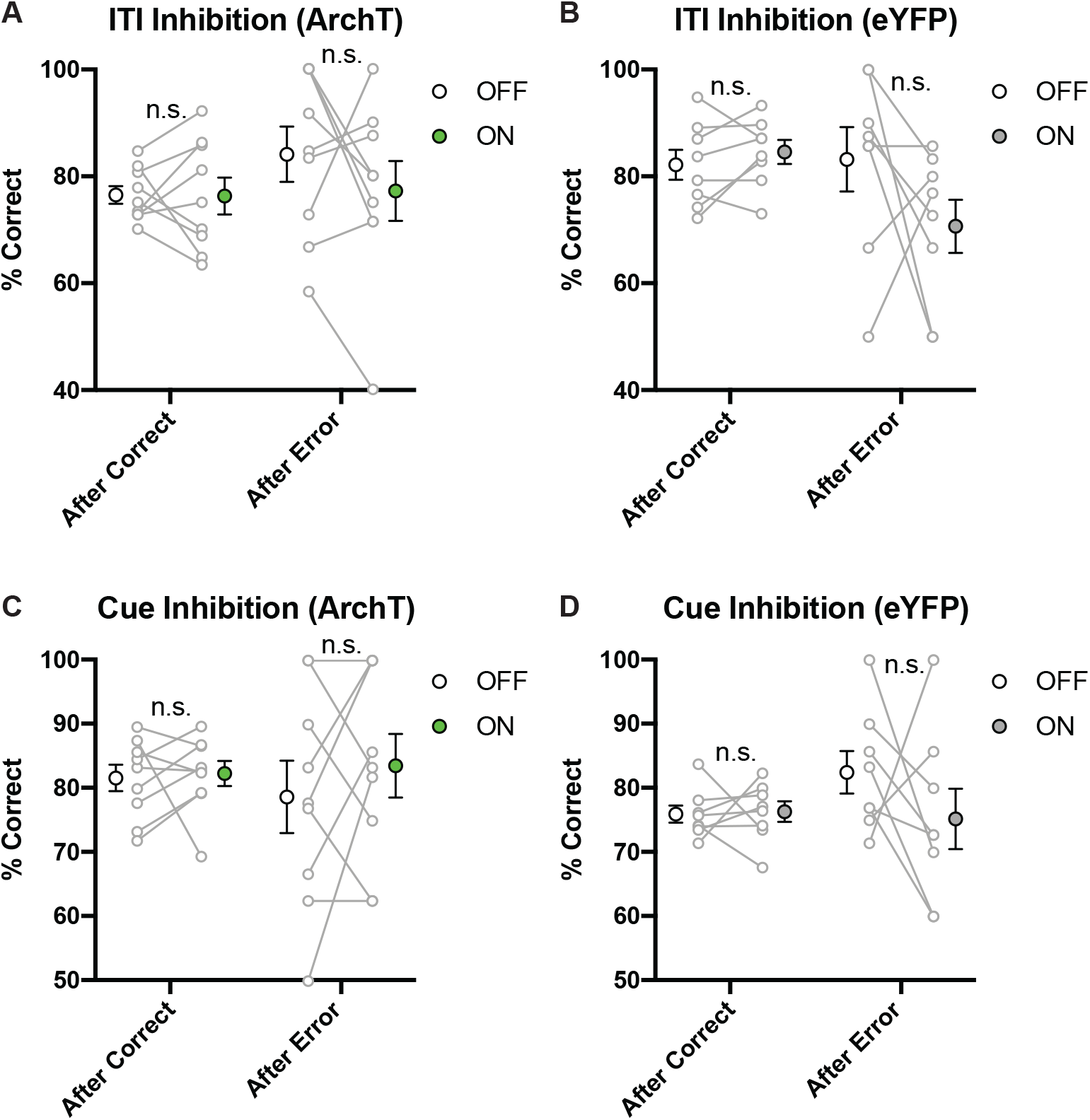
Optogenetic Suppression of D2-MSN in the NAc During ITI or Cue Period Does Not affect the Behavioral Performance in a Recent History Dependent Manner. (A) Optogenetic suppression of D2-MSN in the Nac during ITI period did not affect the behavioral performance after correct or error trial in the VD-Avoid task (Two-way RM-ANOVA with Sidak correction, Treatment effects, F_1,8_ = 1.015, p = 0.3433; After Correct, p = 0.9990; After Error, p = 0.3294). (B) Light stimulation in D2-eYFP during ITI period did not affect the behavioral performance after correct or error trial in the VD-Avoid task (Two-way RM-ANOVA with Sidak correction, Trial type × Treatment interaction, F_1,7_ = 2.535, p = 0.1554; After Correct, p = 0.9265; After Error, p = 0.1910). (C) Optogenetic suppression of D2-MSN in the NAc during Cue period did not affect the behavioral performance after correct or error trial in the VD-Avoid task (Two-way RM-ANOVA with Sidak correction, Treatment effects, F_1,8_ = 0.7479, p = 0.4123; After Correct, p = 0.9760; After Error, p = 0.3471). (D) Light stimulation in D2-eYFP during Cue period did not affect the behavioral performance after correct or error trial in the VD-Avoid task (Two-way RM-ANOVA with Sidak correction, Trial type × Treatment interaction, F_1,7_ = 1.224, p = 0.3051; After Correct, p = 0.9961; After Error, p = 0.3302).

**Supplementary Figure S6.**
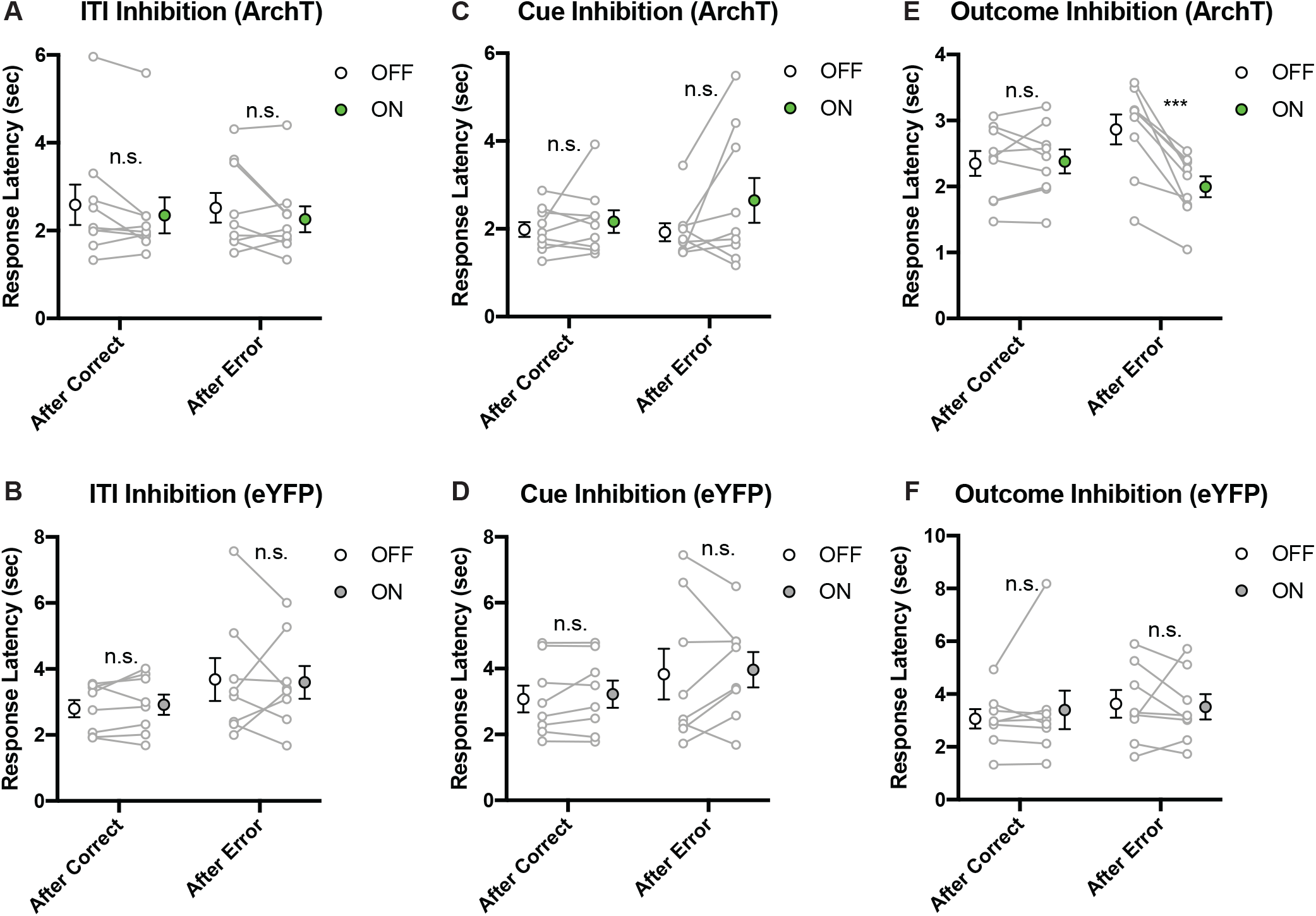
Optogenetic Suppression of D2-MSN in the NAc During Outcome Period After Error Selectively Decreases Response Latency. (A) Optogenetic suppression of D2-MSN in the NAc during ITI period did not affect response latencies after correct or error trial in the VD-Avoid task (Two-way RM-ANOVA with Sidak correction, Treatment effects, F_1,8_ = 0.01032, p = 0.9216; After Correct, p = 0.1918; After Error, p = 0.1553). (B) Light stimulation in D2-eYFP during ITI period did not affect response latencies after correct or error trial in the VD-Avoid task (Two-way RM-ANOVA with Sidak correction, Trial type × Treatment interaction, F_1,7_ = 0.1309, p = 0.7281; After Correct, p = 0.9493; After Error, p = 0.9732). (C) Optogenetic suppression of D2-MSN in the NAc during Cue period did not affect response latencies after correct or error trial in the VD-Avoid task (Two-way RM-ANOVA with Sidak correction, Treatment effects, F_1,8_ = 1.331, p = 0.2820; After Correct, p = 0.8452; After Error, p = 0.1203). (D) Light stimulation in D2-eYFP during Cue period did not affect response latencies after correct or error trial in the VD-Avoid task (Two-way RM-ANOVA with Sidak correction, Trial type × Treatment interaction, F_1,7_ = 0.0003527, p = 0.9855; After Correct, p = 0.9046; After Error, p = 0.9158). (E) Optical stimulation of D2-MSN in the NAc during Outcome period after error response decreased response latencies in the next trial of the VD-Avoid task (Two-way RM-ANOVA with Sidak correction, Treatment effects, F_1,8_ = 23.51, **p = 0.0013; After Correct, p = 0.9716; After Error, ***p = 0.0003). (F) Light stimulation in D2-eYFP during Outcome period did not affect response latencies after correct or error trial in the VD-Avoid task (Two-way RM-ANOVA with Sidak correction, Trial type × Treatment interaction, F_1,7_ = 0.4414, p = 0.5277; After Correct, p = 0.7563; After Error, p = 0.9668).

**Supplementary Figure S7.**
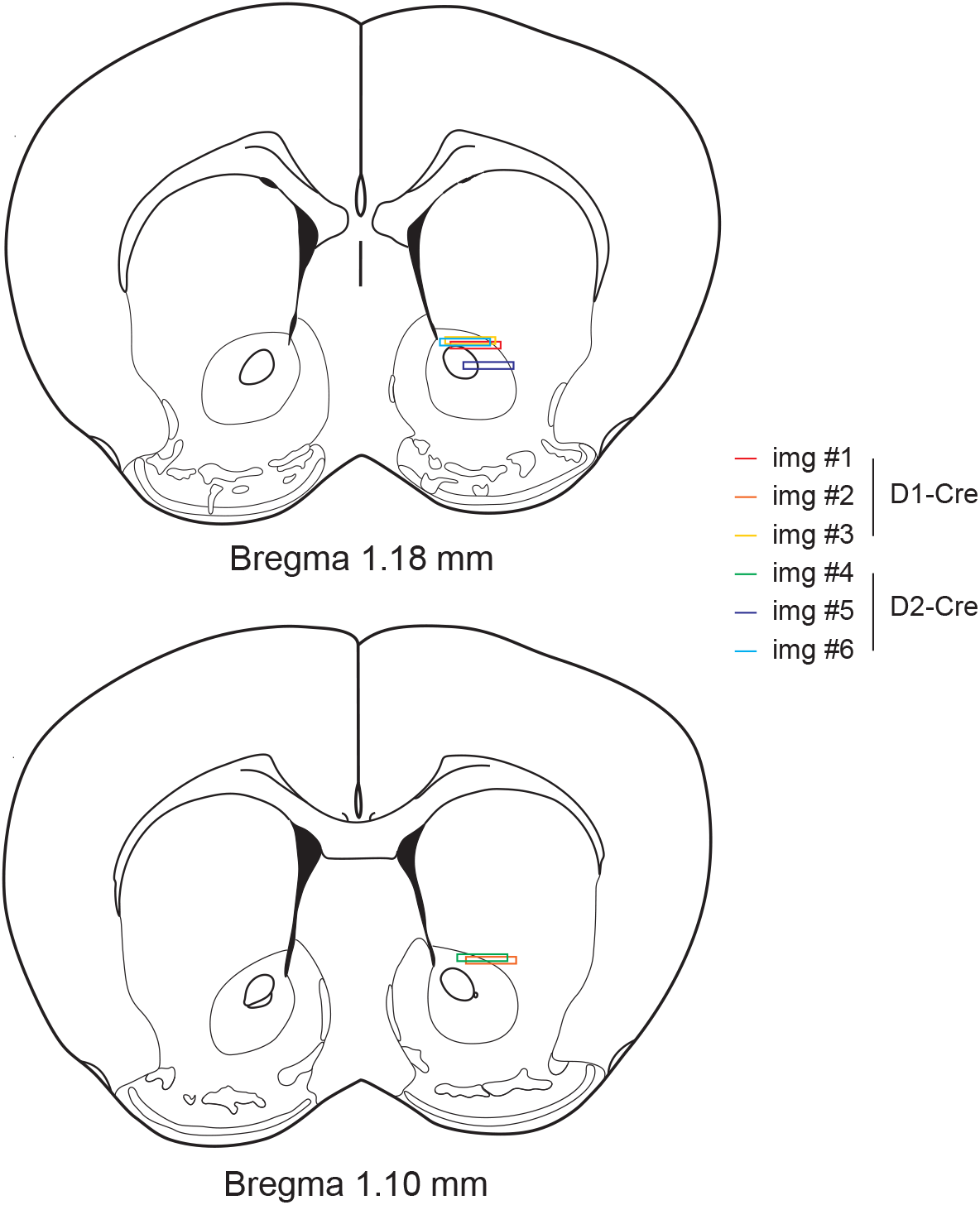
GRIN Lens Placement. Histology on GRIN lens placement.

**Supplementary Figure S8.**
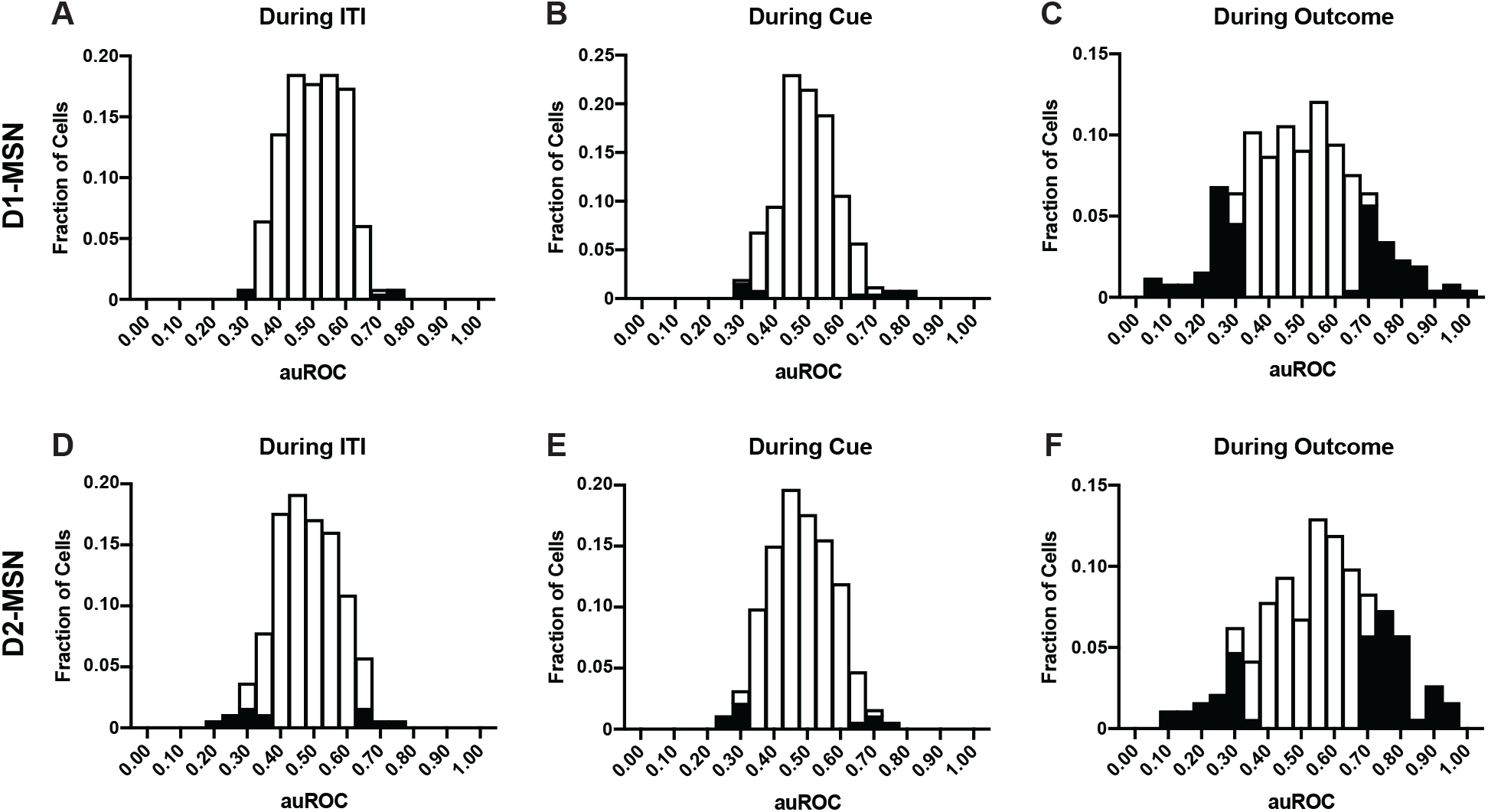
Histograms of auROC During ITI, Cue, and Outcome Period. (A - C) Histogram of averaged auROC of D1-MSN during ITI (A), Cue (B), and Outcome (C) period. (D - F) Histogram of averaged auROC of D2-MSN during ITI (D), Cue (E), and Outcome (F) period.

**Supplementary Figure S9.**
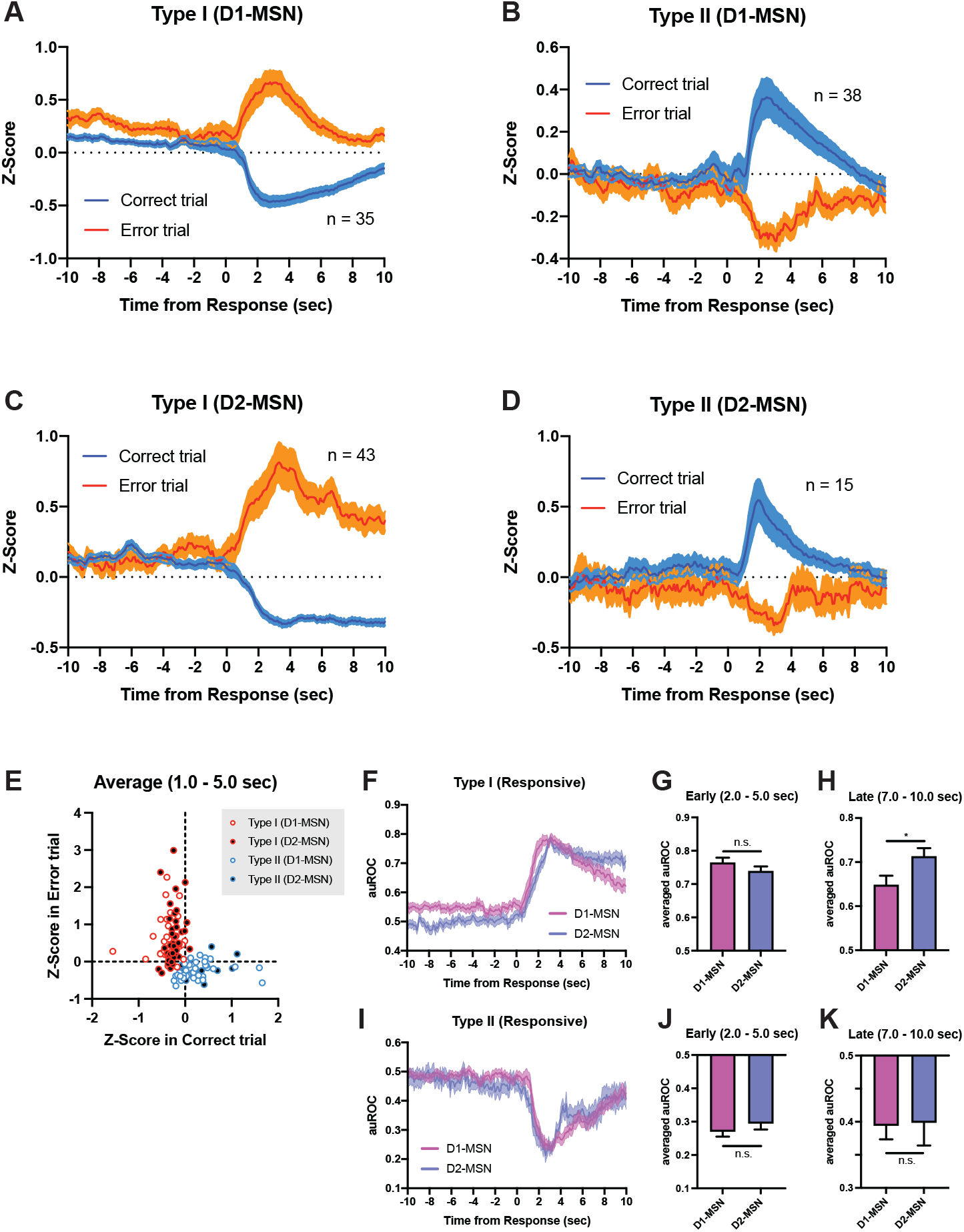
Averaged Calcium Traces from Responsive Type I and Type II Neurons of D1-Cre and D2-Cre mice. (A) Averaged traces of all responsive Type I neurons in D1-Cre mice (n = 35). (B) Averaged traces of all responsive Type II neurons in D1-Cre mice (n = 38). (C) Averaged traces of all responsive Type I neurons in D2-Cre mice (n = 43). (D) Averaged traces of all responsive Type II neurons in D2-Cre mice (n = 15). (E) A scatterplot of individual D1-MSN or D2-MSN responses during Outcome period. (F) Averaged auROC traces of all responsive Type I neurons in D1-Cre and D2-Cre mice. (G) Early responses (2.0 – 5.0 sec) of all responsive Type I (unpaired t-test, t_76_ = 1.289, p = 0.2014). (H) Late responses (7.0 – 10.0 sec) of all responsive Type I (unpaired t-test, t_76_ = 2.373, *p = 0.0202). (I) Averaged auROC traces of all responsive Type II neurons in D1-Cre and D2-Cre mice. (J) Early responses (2.0 – 5.0 sec) of all responsive Type II (unpaired t-test, t_51_ = 0.9313, p = 0.3561). (K) Late responses (7.0 – 10.0 sec) of all responsive Type II (unpaired t-test, t_51_ = 0.1162, p = 0.9080).

**Supplementary Figure S10.**
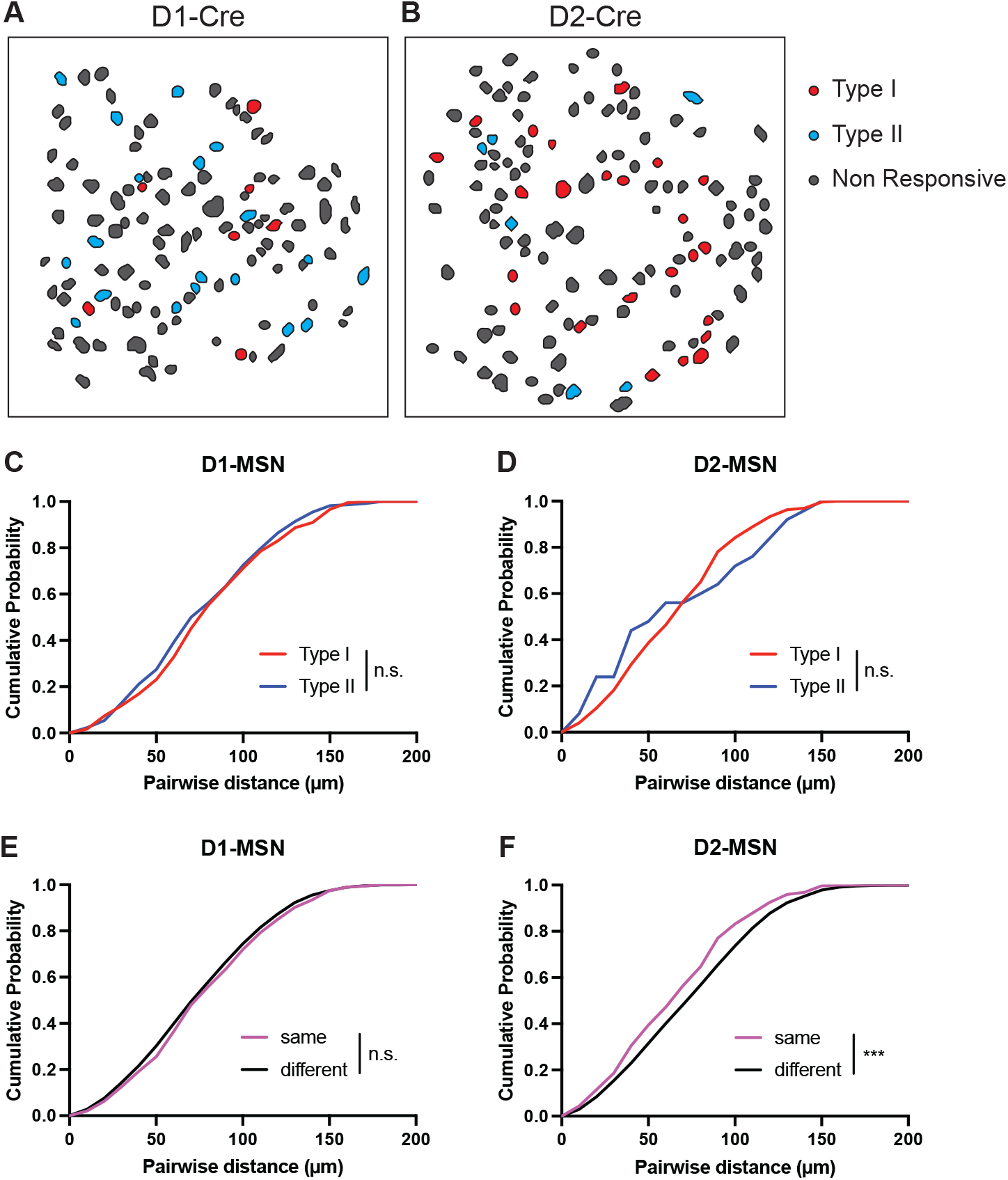
Neurons of the Same Type Stays Closer to Each Other than Neurons of Different Type in D2-MSN. (A and B) Spatial mapping of individual type neurons from representative D1-Cre (A) or D2-Cre (B) mice. (C and D) Quantification of the pairwise distances of each types of neurons in D1-Cre (C, KS test, p = 0.4503) or D2-Cre (D, KS test, p = 0.5357) mice. (E and F) Quantification of the pairwise distances of different types of neurons in D1-Cre (E, KS test, p = 0.2519) or D2-Cre (F, KS test, ***p = 0.0002) mice.

**Supplementary Figure S11.**
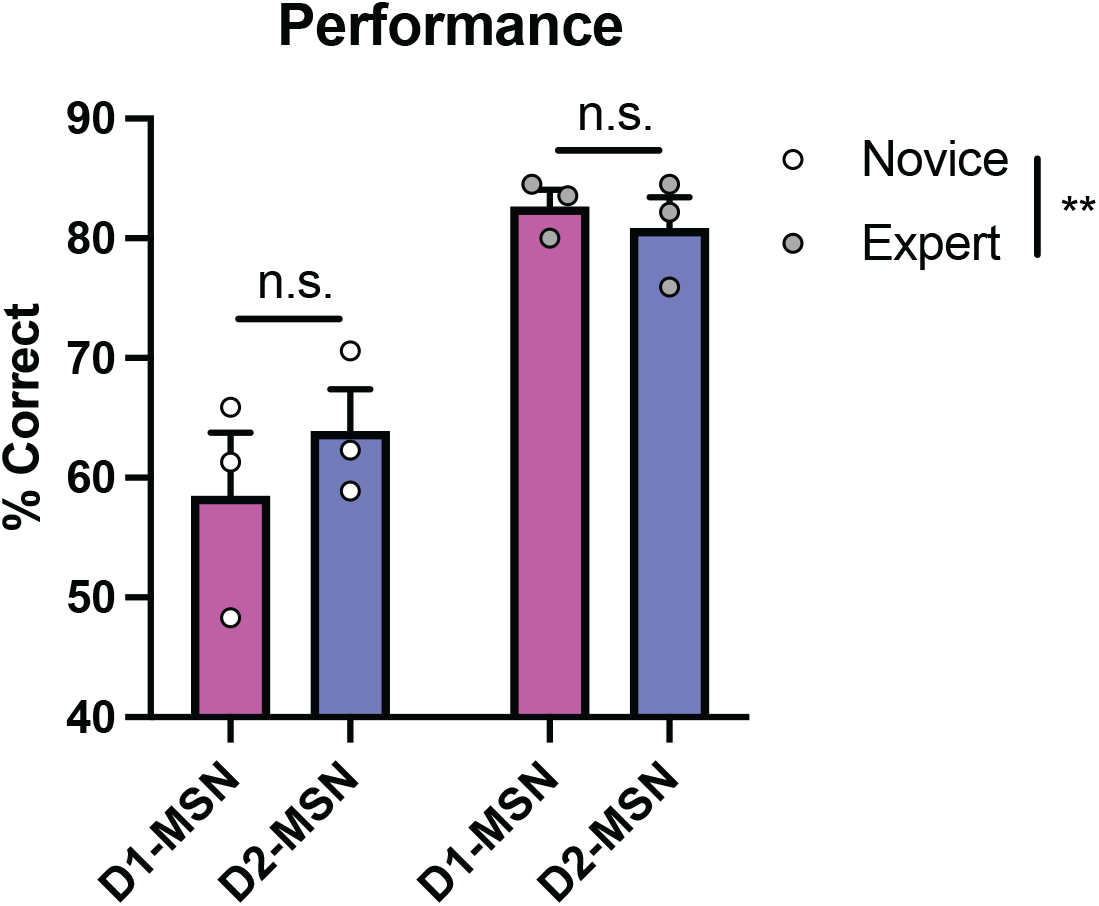
The Behavioral Performance of Novice and Expert Mice. Behavioral performances were improved through learning (Two-way RM-ANOVA with Sidak correction, Learning effects, F_1,4_ = 23.33, **p = 0.0085; Novice, p = 0.3465; Expert, p = 0.8630).

**Supplementary Figure S12.**
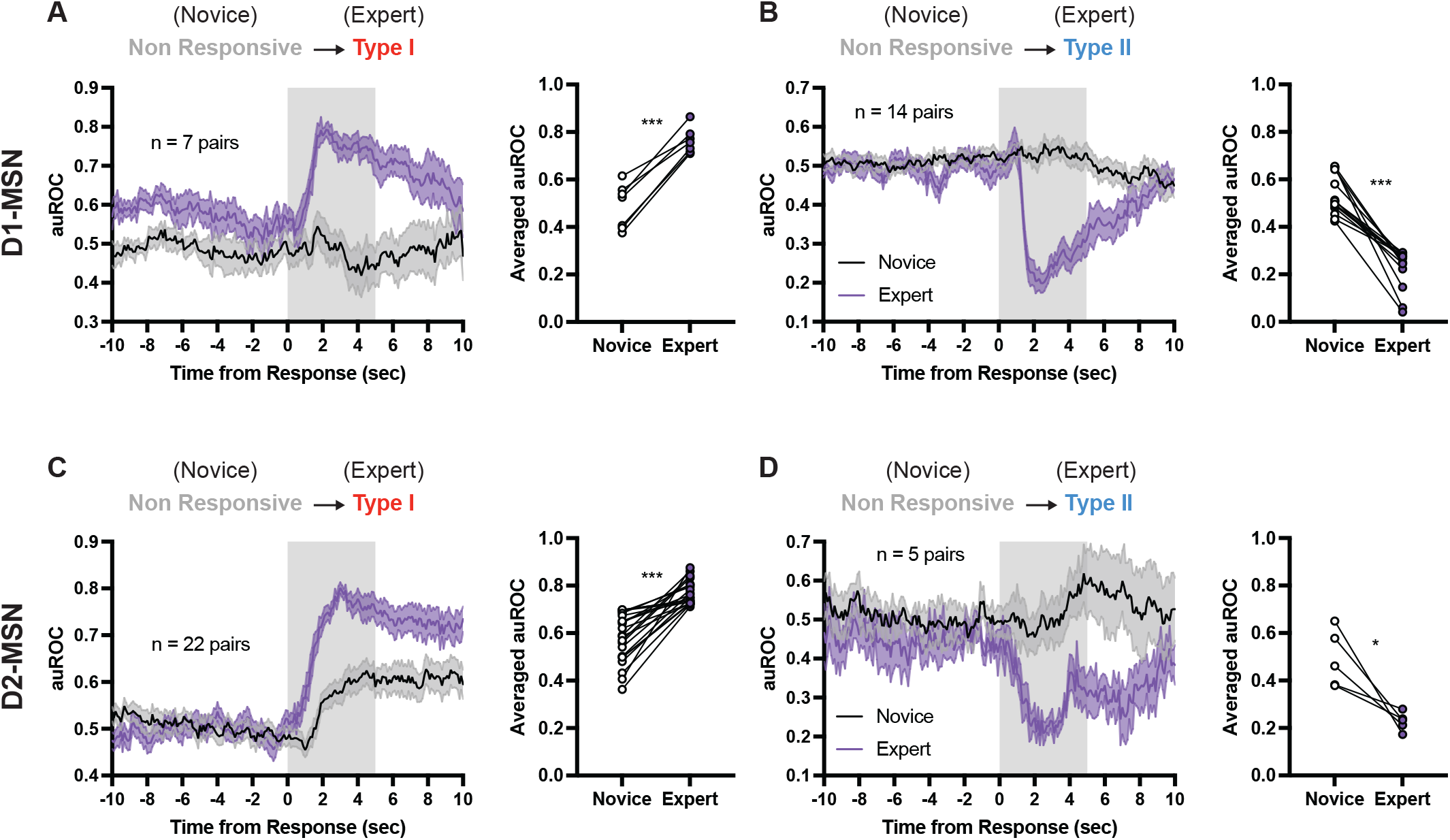
Averaged auROC Traces of Each Type Neuron Through Learning. (A) Averaged auROC traces of neurons becoming responsive Type I from non-responsive type neuron in D1-Cre mice (Left). Averaged auROC in expert mice was significantly higher than that in novice mice (Right, paired-test, t_6_ = 10.22, ***p < 0.0001). (B) Averaged auROC traces of neurons becoming responsive Type II from non-responsive type neuron in D1-Cre mice (Left). Averaged auROC in expert mice was significantly lower than that in novice mice (Right, paired-test, t_13_ = 8.766, ***p < 0.0001). (C) Averaged auROC traces of neurons becoming responsive Type I from non-responsive type neuron in D2-Cre mice (Left). Averaged auROC in expert mice was significantly higher than that in novice mice (Right, paired-test, t_21_ = 7.782, ***p < 0.0001). (D) Averaged auROC traces of neurons becoming responsive Type II from non-responsive type neuron in D2-Cre mice (Left). Averaged auROC in expert mice was significantly lower than that in novice mice (Right, paired-test, t_4_ = 3.841, *p = 0.0184).

**Supplementary Movie 1. Calcium Imaging from Mice Performing the VD-Avoid Task in Correct Trial**

**Supplementary Movie 2. Calcium Imaging from Mice Performing the VD-Avoid Task in Error Trial**

## Notes

### Competing Interest Statement

The authors have declared no competing interest.

